# Experience-Dependent Gain Modulation Drives Thermosensory Responses in Behavior

**DOI:** 10.64898/2026.05.12.724589

**Authors:** Malcom Díaz García, Jonathan Beagan, Ernesto Cabezas-Bou, Matthew J. Thomas, Sandeep Kumar, Lin Shao, Xingyang Fu, Andrea Cuentas-Condori, Jacqueline R. McVay, Josh D. Hawk, Andrew Lauziere, William Mohler, Hari Shroff, Damon Clark, Daniel A. Colón-Ramos

## Abstract

Sensory neurons must extract behaviorally relevant features from dynamic environments while maintaining sensitivity across wide stimulus ranges. To understand how sensory encoding adapts to experience during behavior, we combine long-duration calcium imaging in freely moving *C. elegans* with a temperature-trajectory playback paradigm to determine how the thermosensory neuron AFD extracts behaviorally relevant sensory features during navigation. We observe that AFD functions as a leaky integrator of recently experienced temperature changes, accumulating thermal inputs over a rolling window of tens of seconds, resulting in calcium levels that represent recent temperature dynamics during runs. Importantly, we determine that AFD selectively amplifies responses to temperature changes near its learned preferred temperature. This experience-dependent gain control aligns encoding with the navigational goal, providing a mechanism for representing temperature preference within a derivative-based sensory system. A minimal mathematical model incorporating derivative detection, leaky integration, and temperature-dependent gain captures the calcium dynamics over a range of stimuli, and a simulation based on the mathematical model predicts goal-oriented locomotor strategies across stimulus regimes. Together, these findings show how gain control allows a derivative-based sensory code to represent an absolute goal and guide locomotory strategies during navigation.

**Highlights:**

- AFD encodes recent temperature changes over behaviorally relevant timescales.
- AFD amplifies temperature-change responses within a learned temperature-centered gain window.
- Gain control provides a mechanism for representing distance to temperature preference within the derivative-based sensory system.
- A mathematical model and simulation of derivative sensing, leaky integration, and gain control predicts AFD calcium responses and goal-oriented locomotory strategies across stimulus regimes.

## Introduction

Sensory neurons transform physical stimuli into neural activity that supports perception and behavior. This transformation is not a fixed mapping from stimulus to response. Instead, across sensory modalities, neural responses depend on stimuli history, environmental context, and internal states, reflecting circuit mechanisms that tune sensitivity and dynamic range to extract meaningful features from noisy environments (Bargmann, 2012; Simoncelli and Olshausen, 2001; Smirnakis et al., 1997; Wark et al., 2007). As a result, identical stimuli can evoke different responses depending on the operating regime of the system and the context (Kohn, 2007; Tsukahara et al., 2021). Understanding how sensory representations are shaped by context and experience to extract salient features of the environment and drive behavior remains a central goal in neuroscience.

Thermotaxis in *Caenorhabditis elegans* provides a powerful paradigm to examine how sensory representations support adaptive behaviors. Thermotaxis is a behavior in which animals navigate toward a learned temperature preference based on prior experience (Hedgecock and Russell, 1975). Worms can, for example, remember a cultivation temperature at which they were maintained under favorable conditions for several hours, and, when in a temperature gradient, navigate towards that learned setpoint. This preferred temperature is not innate, and animals can relearn new temperature preferences on the timescale of hours (Hedgecock and Russell, 1975). During thermotaxis, animals must store a representation of the temperature preference, compare the current temperature experience with the temperature goal, and adjust their locomotory strategies to approach, and then maintain, position near the preferred range (Clark et al., 2006; Garrity et al., 2010; Kimata et al., 2012; Luo et al., 2014, 2006; Ryu and Samuel, 2002). While the neural mechanisms that guide thermotaxis are incompletely understood, the core circuitry underpinning this behavior is known and genetically accessible (Hobert et al., 1997; Ikeda et al., 2020; Kimata et al., 2012; Mori and Ohshima, 1995; Satterlee et al., 2001). The possibility of delivering precisely controlled thermal stimuli and visualizing neuronal dynamics as the animals perform a learned behavior makes thermotaxis an attractive model for linking sensory encoding to learned experience and goal-directed navigation.

Within the thermotaxis circuit, the bilaterally symmetric AFD neuron pair serves as the principal thermosensory input (Goodman and Sengupta, 2018). Targeted ablations of AFD disrupt thermotaxis behavior (Chung et al., 2006; Luo et al., 2014; Mori and Ohshima, 1995; Satterlee et al., 2001). Electrophysiological and calcium imaging studies show that AFD responds primarily to changes in temperature rather than absolute temperature, consistent with a derivative-like encoding (Clark et al., 2006; Kimura et al., 2004; Ramot et al., 2008a; Tsukada et al., 2016). Remarkably, AFD detects temperature changes as small as 0.005°C (Luo et al., 2006), yet the neuron supports navigation across thermal landscapes spanning multiple degrees Celsius (orders of magnitude greater than its thermal resolution) during behavior (Clark et al., 2007a; Hedgecock and Russell, 1975; Jurado et al., 2010; Ramot et al., 2008b). The exquisite derivative sensitivity is enabled in part by rapid adaptation mechanisms that preserve AFD’s dynamic range over time (Aoki and Mori, 2015; Goodman and Sengupta, 2018; Hill and Sengupta, 2024; Ramot et al., 2008a). Additionally, the range of temperatures sensed, referred to as operating range, shifts according to recent cultivation history, linking thermosensory physiology to a form of longer-term thermal memory (Kobayashi et al., 2016; Wasserman et al., 2011; Yu et al., 2014). How sensory properties such as rapid adaptation and operating range cohere into a coding strategy that supports thermotactic navigation is not well understood.

To understand which specific features of temperature AFD extracts under varying experiential contexts and how these features support navigation, we established an approach to simultaneously record AFD neuronal activity and behavior in freely moving animals performing thermotaxis. Our system allows us to combine behavioral analysis, naturalistic thermal stimuli, and calcium imaging to investigate which features of temperature AFD extracts and how its encoding properties change with experience. By replaying behaviorally recorded temperature trajectories to immobilized animals, we dissociate sensory encoding from motor feedback and probe AFD responses under precisely controlled yet physiologically relevant conditions. Through these studies, we found that AFD functions as a leaky integrator of recent temperature change (Clark et al., 2006; Tsukada et al., 2016). In this framework, incoming thermal signals are integrated over a time window that progressively discounts older inputs, allowing AFD calcium levels to reflect a temporally weighted history of recent temperature dynamics during forward runs (Abbott and Dayan, 2005). Because AFD is driven primarily by temperature change rather than absolute temperature, this leaky integration gives rise to band-pass filtering, emphasizing behaviorally relevant thermal fluctuations over tens of seconds while attenuating both sustained inputs and rapid noise. Simultaneously, AFD modulates response strength through an experience-dependent gain-control mechanism that multiplicatively amplifies responses to temperature derivatives experienced near the animal’s thermal preference. This gain modulation concept extends prior work showing that AFD has an experience-dependent operating range whose threshold and response window shift with cultivation temperature (Kobayashi et al., 2016; Wasserman et al., 2011; Yu et al., 2014). We formalize these observations in a simple mathematical model in which AFD operates as a derivative-sensing leaky integrator with experience-dependent gain control. By incorporating modeled AFD responses into behavioral simulations, we show how thermosensory signals can guide distinct locomotory strategies during navigation. Together, our experiments and modeling reveal how cultivation temperature-dependent gain control aligns derivative-based sensory encoding with goal-relevant navigation. We believe these findings will generalize, explaining how sensory neurons can use experience-dependent gain control to align derivative-sensing with absolute behavioral goals.

## Results

To investigate how thermosensory information is represented in the nervous system of *C. elegans* during behavior, we developed a system to simultaneously record AFD neuronal activity and behavior in freely moving animals performing thermotaxis (Fig. 1A). We developed a closed-loop tracking and imaging platform that enables long-duration (13-21 minutes) recordings of freely moving *C. elegans* while simultaneously capturing AFD calcium dynamics. Our system builds on earlier freely moving imaging platforms in *C. elegans* that link sensory neuron dynamics to navigation behavior (Ben Arous et al., 2010; Clark et al., 2007a; Faumont et al., 2011; Gengyo-Ando et al., 2017; Jang et al., 2019; Nguyen et al., 2016; Shipley et al., 2014; Tsukada et al., 2016) and extends their capabilities by integrating: 1) an automated tracking system with high spatiotemporal resolution (2.6 μm at 33 FPS;see the Methods section for more information), which enabled sub-second imaging of rapid, cell specific calcium dynamics within freely moving animals; and 2) a machine learning-based analysis pipeline (see Methods section for more information) that detects the animal’s position and extracts ratiometric intensity values of the AFD neuron across time (Fig. 1A, S1). Together, these advances let us collect high-resolution, behaviorally annotated neuronal datasets to quantitatively dissect how AFD encodes temperature-related information and how this encoding aligns with distinct thermotactic strategies.

**Figure 1.**
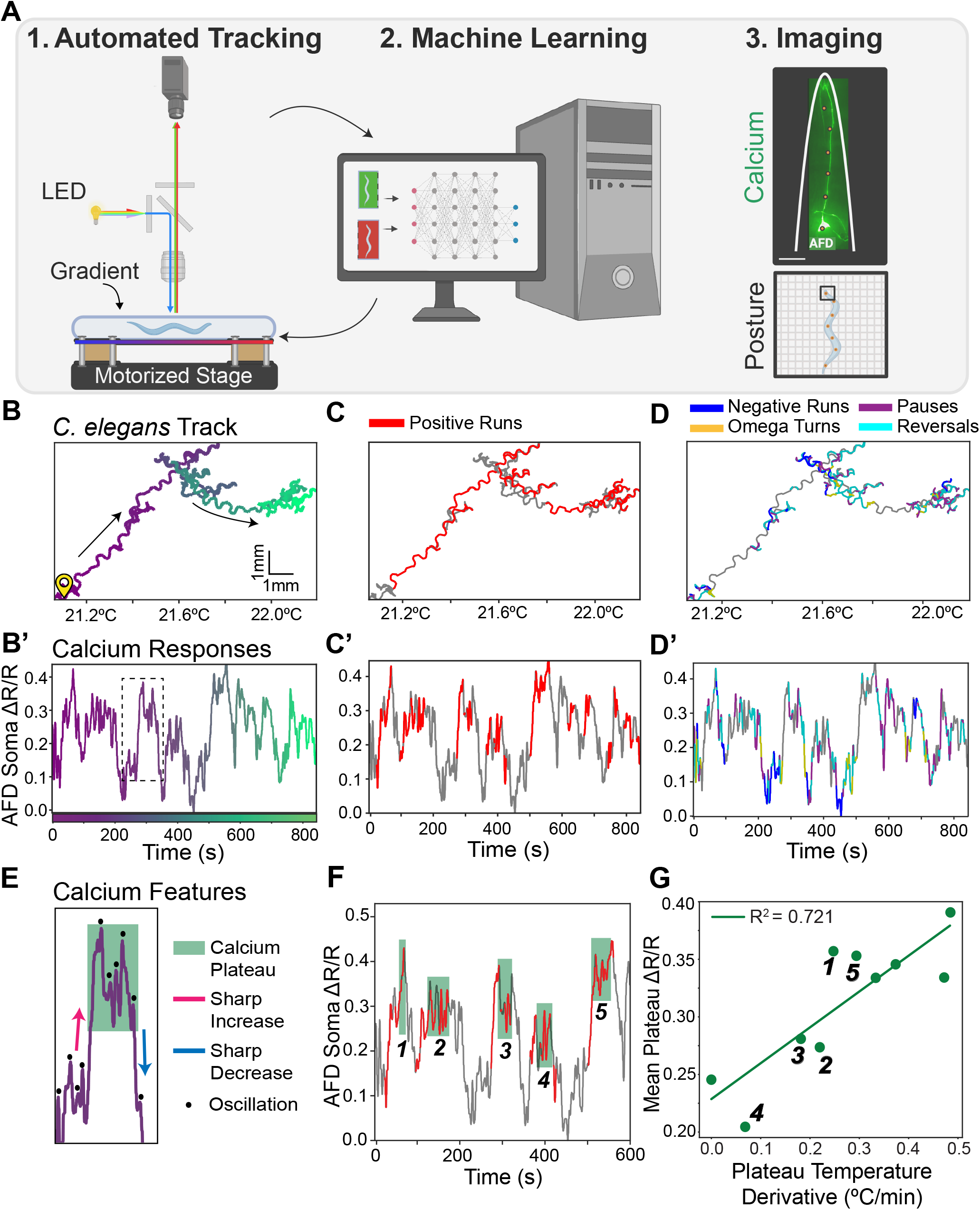
Calcium responses in AFD of freely moving *C. elegans* during thermotaxis behavior. **(A)** Schematic of experimental setup for ratiometric calcium imaging in AFD in freely moving *C. elegans* performing thermotaxis. (1) Animals are placed in a miniaturized thermotaxis assay under a compound microscope, with a motorized stage that is linked to an automatic tracking system. GCaMP6s and RFP are simultaneously imaged. (2) The microscope is connected via a closed-loop system to a computer, which guides the motorized stage to track the animal and keeps it in focus while it is performing thermotaxis behavior. (3) A machine-learning pipeline segments images to extract neuron position and quantify the ratiometric signal from the AFD soma, yielding AFD calcium dynamics to temperature experience (Scale bar, 20 µm.) **(B-D)** Representative thermotaxis trajectory of an animal trained to prefer 25°C (Tc25). **(B)** Track color-coded by elapsed time (purple to green) and **(B’)** AFD calcium responses (ΔR/R) corresponding to the same timestamps as the track in **(B)**. Black arrows in **(B)** point to net movement direction of the animal, and dashed box in **(B’)** corresponds to zoomed-in image in **(E). (C)** Track from **(B)**, with positive runs (sustained movement towards a warmer temperature) colored red. **(C’)** Calcium traces, as in **(B’)**, but colored red for run events corresponding to **(C). (D-D’)** As **(C-C’)**, but annotated for reversals, omega turns, pauses and runs towards a colder temperature (negative runs). **(E)** Zoomed-in image corresponding to dashed box in **(B’)**, highlighting the features observed in the calcium responses, including large increases and decreases in AFD activity (red and blue arrows), calcium plateaus (shaded region) and oscillations (black dots). **(F)** As in (B’), but with calcium plateaus highlighted and numbered. **(G)** Relationship between mean AFD Ca^2+^ and mean temperature derivative across each plateau event (numbered points correspond to the plateaus in **(F)**), with a linear fit between mean plateau value of ΔR/R and plateau temperature derivative (°C/min). Variance to fit in plateau height (R^2^ = 0.72).

We raised animals in conditions that elicit positive thermotaxis behaviors, defined as movement from a cooler starting temperature up a thermal gradient toward their preferred cultivation temperature (Hedgecock and Russell, 1975; Mori and Ohshima, 1995), and recorded AFD calcium activity and animal position and posture as they freely navigated the gradient. As they explored the gradient for several minutes, the animals traversed ~1°C using a movement strategy that resulted in net migration up the gradient toward their cultivation temperature (Fig. 1B). AFD is known to produce graded calcium responses to warming, and consistent with this, we observed that AFD activity was modulated during thermophilic behavior (Fig. 1B’). Comparison of the temperature trace with AFD calcium dynamics revealed that AFD activity did not simply track absolute temperature, nor did it monotonically increase with warming (Fig. S2D). Instead, AFD responses were constrained within a bounded range of ΔR/R values, corresponding to experienced temperature derivatives between 0.1-0.6°C/min range (Fig. 1B’, S2D). Within this range, we identified three distinct patterns of AFD calcium activity (Fig. 1B’ and Fig. 1E): 1) Large, transient increases and decreases in calcium levels, often appearing as sharp deflections (arrows in Fig. 1E); 2) Sustained ‘calcium plateaus’, where AFD activity stabilized at an elevated or intermediate level for tens of seconds (green shades in Fig. 1E and Fig. S2C’); and 3) Small, periodic oscillations in AFD calcium activity, persistently observed across the gradient and independent of the underlying baseline calcium level (highlighted with dots in Fig. 1E).

To understand how these calcium dynamics relate to behavior, we aligned behavioral annotations with the AFD activity patterns (Fig. 1C-D). We first investigated how the larger calcium transients relate to behavioral state transitions by annotating the distinct strategies employed, including positive and negative runs, omega turns, reversals and local searches. We then related the timing of distinct behavioral motifs to the AFD calcium traces (Figs. 1C-D and S2A-B). We found that sharp increases in calcium levels were closely associated with the initiation of thermophilic runs, during which animals actively migrated up the temperature gradient (Fig. 1C-C’ and S2A-A’). Conversely, these runs frequently terminated with sequentially executed negative runs, turning behaviors (omega turns and reversals) and local searches, which coincided with large decreases in AFD activity (Fig 1D-D’ and S2B-B’). These observations suggest that large changes in AFD calcium levels correlate with transitions between navigational states. Specifically, thermophilic runs appear to be “bookended”by opposing calcium shifts: a sharp rise in AFD activity at the onset of a thermophilic run, and a sharp decline at the transition into a sustained series of reorientation behaviors that shifts the heading of the organism.

While large AFD calcium increases often marked the initiation of thermophilic runs, run persistence resulted in a sustained plateau in calcium activity (green shaded regions in Fig. 1E-F and S2C’) that lasted the remainder of the run (for tens of seconds) (Fig. S2E). These “plateau phases”provided an opportunity to examine how AFD Ca^2+^ levels relate to ongoing thermal experience during forward navigation. To assess this, we plotted the average AFD calcium signal during plateau periods against the average rate of temperature change (ΔT/Δt) experienced during those same intervals. These analyses revealed a strong positive correlation (R^2^=0.72) between plateau average calcium levels and ΔT/Δt (Fig. 1F-G). In contrast, when we examined the correlation between instantaneous ΔT/Δt and calcium activity across all time points, we found no clear relationship between the average AFD calcium signal and the instantaneous rate of temperature change (ΔT/Δt) (Fig. S2F). Our findings are consistent with previous studies which showed that AFD encodes temperature derivatives (Clark et al., 2006; Kimura et al., 2004; Ramot et al., 2008a; Tsukada et al., 2016), and further support that the AFD calcium levels reflect, not moment-to-moment readouts of instantaneous temperature changes, but an integrated measure of thermal inputs over time.

The interpretation that AFD responses reflect a temporally weighted history of sensory inputs is consistent with prior work showing that a temperature upstep evokes a transient AFD response that adapts to pre-stimulus levels on the timescales of ~40 seconds (Clark et al., 2007a). It is also consistent with studies that estimated the AFD response function during thermotaxis and found that AFD integrates recent temperature changes over a finite temporal window of approximately 20 seconds (Tsukada et al., 2016). To further examine this in our system, we also derived the response function of AFD by linking recent ΔT/Δt history to current AFD activity for the calcium dynamics obtained from our freely moving animals. Using least-squares estimation (see Methods), we fit the response function to a standard exponential decay function (Fig. S2G). While our sensors and assays are distinct from those used in previous studies, we observed that the general features of the response function were consistent with those previously reported (Tsukada et al., 2016), displaying temporal integration that extended at the tens of seconds timescale, with the influence of prior stimuli on the current response exponentially decreasing with time. We note that the tau of the response function derived from our studies (~50 seconds) is also similar to the ~45s tau of the adaptation kinetics for the AFD sensory thresholds (Hawk et al., 2018). Our findings support a model in which AFD acts as a leaky integrator of recent temperature-change signals. By summing temperature-change signals over time while allowing older inputs to decay, this computation produces a low-pass-filtered representation of recent thermal experience with a tens of seconds-scale exponential time constant. Because this filtering operates on temperature-change signals rather than absolute temperature, the resulting response is band-pass-like with respect to the original temperature trajectory, attenuating both sustained temperature and high-frequency temperature fluctuations. Together, our findings support that ongoing calcium activity in AFD reflects a temporally weighted history of temperature dynamics, suppressing rapid fluctuations while preserving behaviorally relevant thermal trends.

Based on the findings that AFD acts as a leaky integrator, we hypothesized that the observed plateau in the calcium dynamics of freely moving animals might reflect the temperature derivative experienced during the thermophilic run interval, filtered by the neuron’s intrinsic decay kinetics. To causally test this, we used a Peltier device capable of delivering precise, controlled thermal stimuli to immobilized animals (Hawk et al., 2018). This system allowed us to systematically vary the rate of temperature change (ΔT/Δt), enabling direct testing of the hypotheses derived from our freely moving recordings. We applied three protocols in which we independently varied components of the temperature derivative, including increasing temperature differences over a fixed amount of time (first protocol), a fixed change in temperature over increasing durations (second protocol), or an increase in the temperature derivative and the absolute temperature (third protocol).

In the first protocol, we tested whether increasing temperature derivatives via higher ΔT over fixed amounts of time led to a proportionally higher AFD calcium level. To achieve this, animals were exposed to five 1-minute linear warming ramps, with incremental increases of temperature steps ranging from 0.2°C to 1.0°C (in the range of 20°C to 21°C in absolute temperatures) interleaved with 5-minute holds at baseline temperature (20°C). We observed that AFD calcium responses increased in magnitude with ramp steepness, showing a strong positive correlation between the rate of temperature change and the peak calcium signal (Pearson coefficient of 0.92), up to approximately 0.8–1.0°C/min (Fig. 2A and Fig. S3A). This relationship was linear in the range of 0.2°C/min to 0.6°C/min, after which AFD’s ability to represent a temperature derivative appeared saturated (Fig 2A and Fig. S3A). Notably, the linear range of derivative representation observed in the controlled thermal stimuli (~0.1–0.6°C/min) matched the range of derivatives sampled during positive runs by freely moving animals (Fig. 1G). These results suggest that AFD operates within an approximately linear encoding range for temperature derivatives, and that animals may modulate their navigational strategies to remain within this sensitive range. Importantly, our data indicates that the sustained calcium plateaus observed in freely moving animals (Fig. 1F-G and S2C’) correspond to AFD’s integrated, band-pass-filtered encodings of temperature derivatives during thermophilic runs.

**Figure 2.**
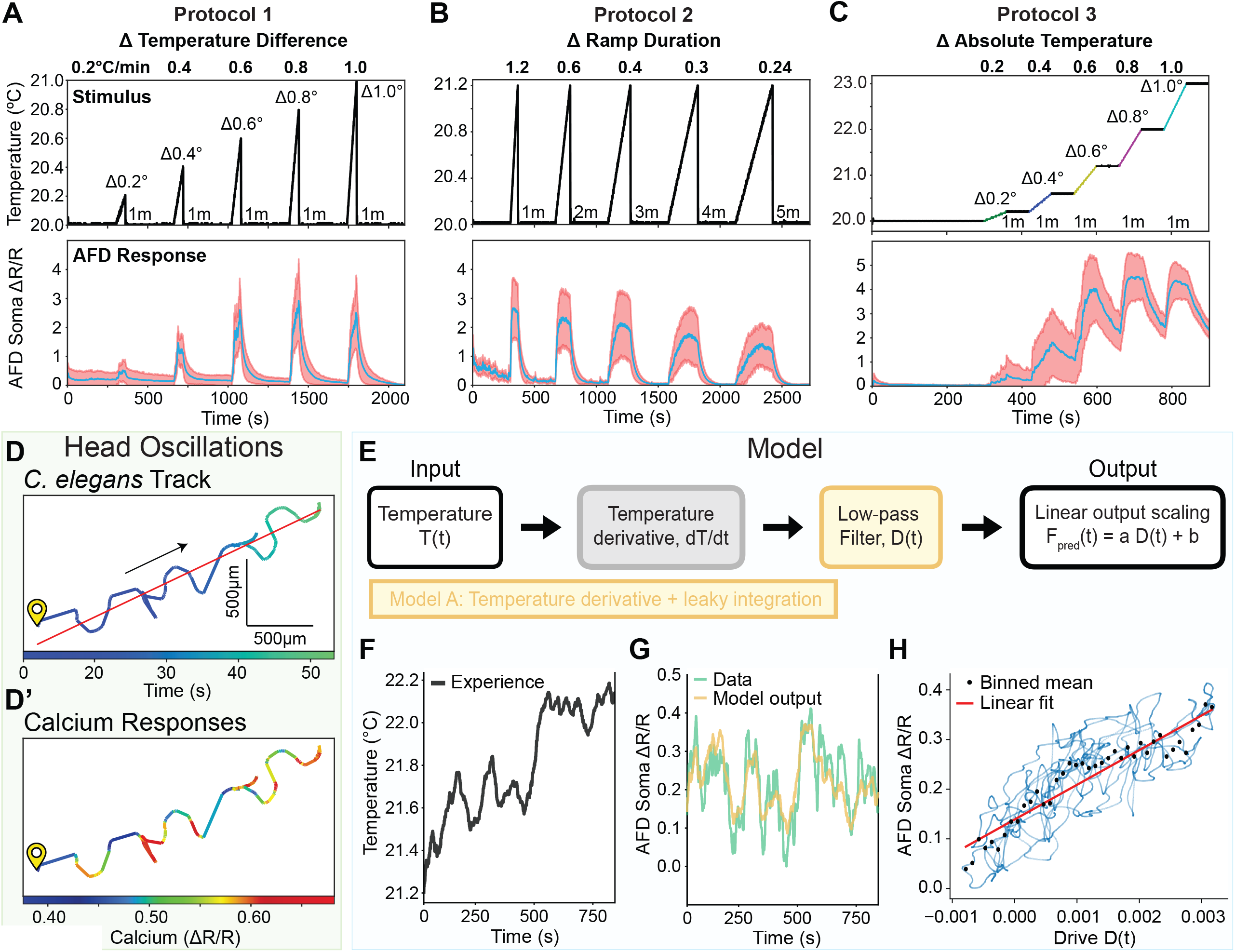
Relationship between sustained AFD calcium responses and temperature derivatives. **(A)** AFD Ca^2+^ responses (bottom panel, n=14 animals) to heating ramps (top panel) with resets to baseline temperature (20°C) between ramps. Ramp duration was fixed with increasing maximum temperature (increasing ΔT, hence larger temperature derivatives). Blue trace in bottom panel denotes mean with 1 standard deviation shades in red. Each ramp was labeled with its corresponding ΔT at the peak of the ramp. **(B)** As in (A), but Ca^2+^ responses (n=8 animals) to heating ramps with maximum temperature fixed with increasing ramp duration (increasing Δt, hence smaller derivatives). **(C)** As in (A), but Ca^2+^ responses (n=15 animals) to heating ramps without resetting to baseline between each ramp (increasing ΔT without baseline resets), to examine the influence of absolute temperature while maintaining the same sequence of temperature derivatives. Each ramp was labeled with its corresponding ΔT at the peak of the ramp. (D) Behavioral track of a representative plateau event from the freely moving thermotaxis experiment shown in Fig. 1, denoting head oscillations resulting from lateral bending. The blue to green gradient (top) indicates elapsed time across the positive run. **(D’)** Same track shown in **(D)**, but colored with a rainbow gradient according to AFD Ca^2+^ level (blue lowest and red highest), highlighting the stereotyped oscillations in signal as the worm undulates. (E) Schematic of minimal mathematical model linking temperature experience to AFD calcium responses. Experienced temperatures are used to derive rates of change 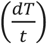. Then, a low-pass filter generates a latent variable *D*(*t*), modeling the leaky integration property. Lastly, a Ca^2+^ prediction *F*_*pred*_(*t*) is obtained by linearly scaling *D*(*t*). **(F)**. Thermal experience over time recorded from the freely moving animal shown in Fig. 1. **(G)** For the same experiment in **(F)**, the mean AFD soma Ca^2+^ (ΔR/R) (light green trace) is shown with the model prediction *F*_*pred*_(*t*) in yellow. **(H)** Binned average measured Ca^2+^ (ΔR/R) against model derived *D*(*t*). Black points denote binned means of the individual time points in blue, and the red line is a linear fit.

Next, we asked whether the same relationship was held when the total temperature change (ΔT) was fixed, but the duration (Δt) of each ramp varied. In this second protocol, animals experienced five consecutive ramps with a constant 1.2°C increase from baseline, but with ramp durations ranging from 1 to 5 minutes, resulting in the same absolute temperature being achieved, but via progressively shallower slopes (1.2°C/min to 0.24°C/min). AFD calcium levels again scaled with ramp steepness: slower ramps produced lower peak calcium signals (Fig. 2B). All ramps reached the same final temperature, confirming that the graded responses were driven by ΔT/Δt and not by the absolute temperatures achieved during the gradient. A positive correlation (Pearson coefficient of 0.89) between ΔT/Δt and peak calcium levels was observed again in this protocol, extending across the 0.1–0.6°C/min range (Fig. S3B) and consistent with the linear range of AFD observed for the first protocol and for freely moving animals. This finding further reinforces the relationship between AFD calcium levels and experienced ΔT/Δt, supporting a model in which the calcium “plateaus”observed during thermophilic runs in freely moving animals reflect the temperature derivatives experienced by the animal during those runs (Fig 1F and S2C’).

In the third protocol, we tested how AFD encodes the temperature derivative as the animal’s absolute temperature exposure increased. We tested this by not returning the absolute temperature to the same baseline after each ramp. Instead, each 1-minute ramp increased the peak temperature by 0.2°C (0.2–1.0°C/min), and the new peak was maintained as the baseline for the following ramp, resulting in a cumulative increase from 20°C to 23°C, similar to what an animal migrating on that temperature range might experience. We observed that AFD calcium responses again scaled with ΔT/Δt, producing graded responses that rose with ramp steepness (Fig. 2C). Moreover, a linear fit of peak responses versus ΔT/Δt showed a strong positive correlation across the 0.1–0.6°C/min range (Fig. S3C). Taken together, these results reveal that AFD dynamically tracks temperature change by generating graded calcium responses and sustained plateaus that reflect both the magnitude and persistence of the temperature derivative stimulus. These dynamics observed in freely moving animals reflect the properties of AFD as a leaky integrator, its calcium levels representing a short-term (tens of seconds) temperature derivative history that stabilizes as animals perform a run in a temperature derivative of a relatively constant value.

Even for the periods which we termed “plateaus”in freely moving animals (Fig. 1F and S2C’), AFD responses displayed small, rhythmic oscillations in calcium levels which were not observed for immobilized animals stimulated with a linear ramp (Compare Fig. 1E-F and S2C’ to Fig. 2A-C). We analyzed these oscillations and asked whether they exhibited consistent frequency and phase relationships that aligned with a navigational strategy of the animal. We found that the pattern and frequency of the calcium oscillations aligned with the undulating head bends of the freely moving animals (Fig. 2D-D’). Specifically, when the frequency and phase of the calcium responses and head oscillations were compared (See Methods), we observed a negligible frequency difference (0.005 Hz difference between the head oscillation and the calcium oscillation) and phase lag (4π/11) between head bending and the oscillating calcium responses (Fig. S3D-D’’). Our values were similar to prior measurements of calcium oscillations in AFDs of partially immobilized and freely moving animals (Clark et al., 2007a). We then examined if the calcium oscillation amplitudes corresponding to the head bends held a relationship to the magnitudes of the derivatives of rhythmic oscillations in the plateaus. We quantified the amplitude of the oscillatory responses for plateaus corresponding to varying derivative values and observed that the amplitude of the oscillatory responses remained relatively similar regardless of the calcium plateau level and derivative (Fig. S2H-I). These results suggest that AFD is capable of sensory encoding that allows this sensory neuron to track tonic changes in ΔT/Δt during prolonged runs up a gradient while simultaneously remaining sensitive to rapid, small thermal fluctuations. This property of AFD allows it to encode accurate temperature derivatives of varying magnitudes across diverse gradient conditions while simultaneously being able to sensitively detect small temperature fluctuations, like those experienced during undulatory head-bends as the animals navigate a temperature isotherm. These patterns highlight how AFD calcium dynamics encode temperature information across multiple behavioral scales, from fine-grained oscillations that reflect local temperature fluctuations due to head-bends, to large-amplitude changes that correspond to transitions in run strategies.

Having identified roles for leaky integration and derivative detection in AFD calcium dynamics, we next developed a minimal mathematical model to formalize these properties and test their relative contributions to temperature encoding (Fig. 2E). The temperature derivative, 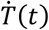, is the experienced temperature, T(t), converted into its instantaneous rate of change, 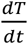, an approximation of a derivative-taking computation that likely occurs over seconds (Clark et al., 2006). This derivative is then passed through a low-pass filter to generate a latent drive variable, D(t), which represents the output and captures temporally-integrated temperature changes. This low-pass filtering is given by the equation:

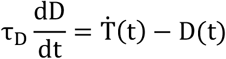

where τ_D_ is the timescale over which the system integrates changes in temperature. A linear scaling step transforms this drive into a predicted calcium response:

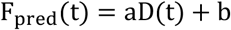

Here, the constants a and b linearly map the latent drive to the measured fluorescence, accounting for differences in signal scaling and baseline across recordings. When applied to temperature stimuli experienced by freely moving animals navigating towards their preferred cultivation temperature (Fig. 2F), this model captured the dominant temporal dynamics of experimentally observed AFD calcium responses (Fig. 2G), including transient increases during warming, decay during cooling, and sustained calcium plateaus. We then evaluated performance within a linear-nonlinear (LN) framework. In a linear-nonlinear (LN) framework, a linear filter extracts specific features from a stimulus, which are then transformed by a static nonlinearity to predict the response (Chichilnisky, 2001; Schwartz et al., 2006). To evaluate our model in this context, we plotted measured calcium responses as a function of the model-derived drive, D(t) (Fig. 2H). The blue points show the raw relationship between D(t) and calcium responses, while the black circles represent binned means obtained by averaging responses within bins of D(t). The binned averages revealed a predominantly linear relationship (Fig. 2H), indicating that when freely moving animals navigate near their cultivation temperature, AFD responses can be approximated as a linear transformation of low-pass filtered temperature derivatives. Thus, under restricted stimulus conditions centered around the preferred temperature, AFD dynamics are well described by a derivative-based linear model that incorporates leaky integration.

Our use of a linear model to generate a minimal dynamical formulation of AFD responses builds on and extends prior response-function analyses of AFD temporal filtering (Clark et al., 2007a; Tsukada et al., 2016). This formalization allows us to separately quantify the contributions from variables such as derivative sensitivity and temporal integration, and to consider additional, context-specific variables that might also shape the AFD response dynamics. We therefore used this framework to revisit a longstanding observation in the field: that AFD’s response range shifts with cultivation history. These shifts have often been interpreted as a form of temperature memory stored within AFD for three reasons: 1) they occur on timescales comparable to changes in behavioral preference (hours); 2) they parallel changes in behavioral preference; and 3) they define the temperature range over which AFD responds in relation to the cultivation temperature experience (Kobayashi et al., 2016; Wasserman et al., 2011; Yu et al., 2014). However, it has also been shown that AFD response threshold can adapt within minutes (Hill and Sengupta, 2024; Ramot et al., 2008a; Yu et al., 2014), reflecting recent thermal experience rather than long-term cultivation temperature (Hawk et al., 2018), which is happening in similar timescales as the AFD’s leaky integration of temperature derivatives. We therefore asked how previous thermal experience, including recently experienced temperature and longer-term cultivation-temperature memory, shapes AFD responsiveness.

We first examined how longer-term thermal experience shapes AFD encoding within the framework of a short-term, derivative-sensitive leaky integrator. We achieved this by repeating and extending cultivation-temperature shift paradigms across multiple timescales (Hawk et al., 2018; Kobayashi et al., 2016), allowing us to examine AFD responses under varying thermal histories while accounting for fast-adaptive threshold dynamics. Specifically, we exposed animals cultivated at 15°C (Tc15) to 25°C for varying durations, including the standard 4-hour exposure period resulting in the temperature preference memory as well as shorter periods on minutes timescales known to affect the temperature threshold, but not the temperature preference (Hawk et al., 2018) (Fig. 3A-A’). We also performed the converse experiment, cultivating animals at 25°C (Tc25) and exposing them to 15°C for varying durations (Fig. 3B). We then presented a long-lasting sinusoidal temperature ramp spanning much of the physiological range of *C. elegans* (12–28°C) (Fig. 3). When Tc15 animals were exposed to 25°C for minutes, and then presented with a temperature ramp, the temperature at which AFD first responded (the AFD response threshold) shifted to the warmer-experienced temperatures within minutes (Fig. 3A, black arrow). These findings are consistent with rapid adaptation of the AFD response threshold (Hawk et al., 2018). Under these conditions of exposing Tc15 animals to 25°C for minutes, we did not observe a translocation of the response profile of AFD (Fig. 3A, star). Over longer exposures to 25°C, we observed that the entire response profile shifted toward the new cultivation temperature (Fig. 3A’, red arrow), consistent with previous observations of the operating range (Kobayashi et al., 2016; Wasserman et al., 2011; Yu et al., 2014). Similar results were observed when animals were cultivated at 25°C (Tc25) and shifted to 15°C for varying durations (Fig. 3B-B’’). Together, our experiments disentangle the contributions of cultivation-dependent response shifts and rapid threshold adaptation, revealing how distinct forms of plasticity jointly define AFD’s operating range.

**Figure 3.**
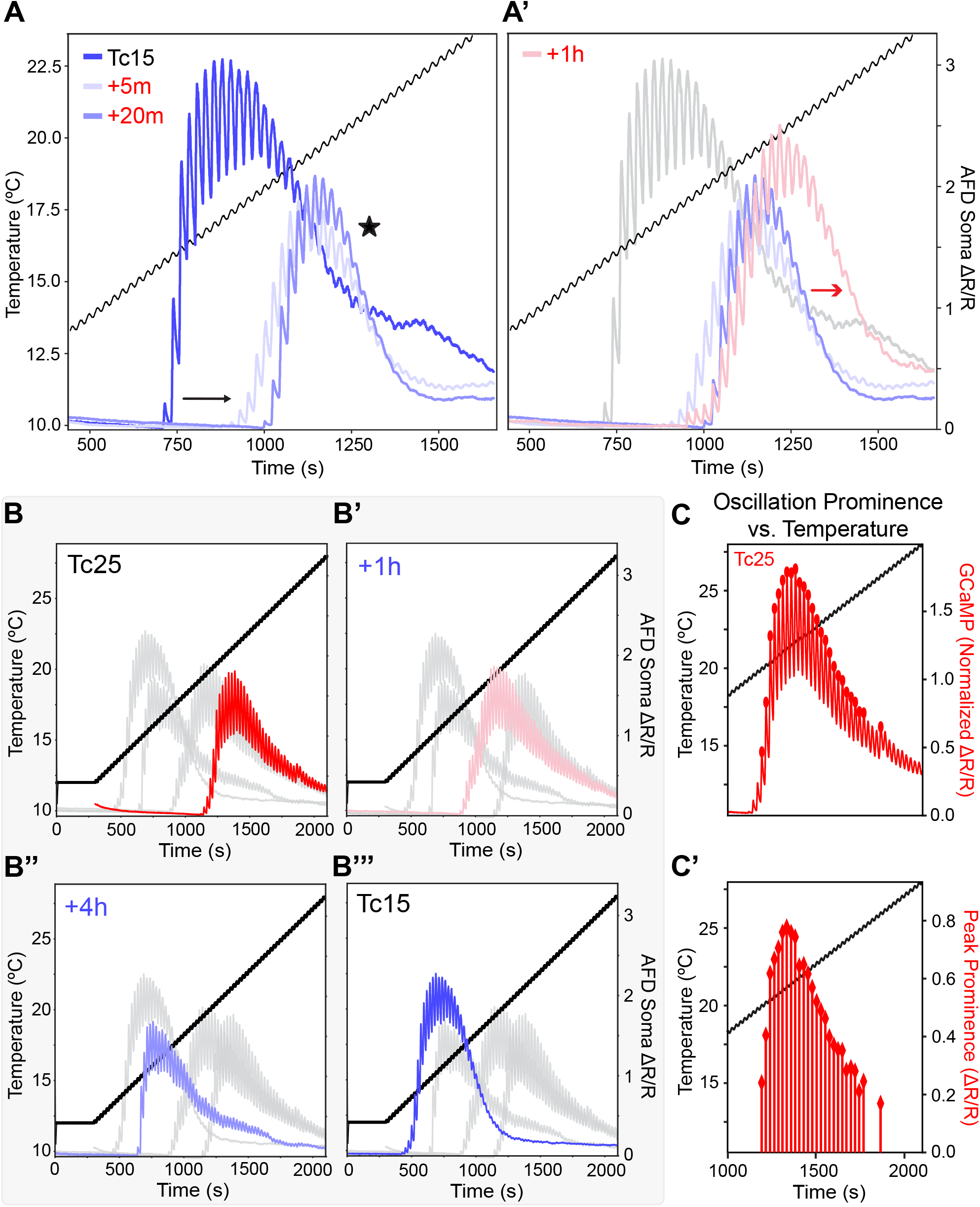
AFD temperature threshold and gain region adapt on two distinct timescales to temperature changes. **(A-B)** AFD responses to a sinusoidally oscillating stimulus overlaid on a 0.5°C/min temperature ramp (black lines). **(A)** Three AFD responses, each curve representing the mean response across multiple animals. Responses from Tc15 (n=5 animals) trained (no additional temperature exposure) worms are colored blue;Tc15 (n=10 animals) worms with 5min hold at 25°C are colored light blue;Tc15 (n=12 animals) worms with 20min hold at 25°C are colored periwinkle. After these brief exposure intervals, the initial response threshold of AFD to the stimulus shifts (denoted by black arrow), while sensitivity at higher temperatures does not substantially change (denoted by the star symbol). **(A’)** The three responses from **(A)** but adding Tc15 worms (n=10 animals) with 1hr hold at 25°C highlighted in pink. Note how, only in the 1h hold condition, the range of sensitivity at higher temperatures increases (red arrow). **(B-B’’’)** Four AFD responses to temperature downshifts of different durations. Each curve represents the average response across all animals. **(B)** Highlighting Tc25°C trained worms in red (no additional temperature exposure) (n=17 animals, 2 independent experiments); **(B’)** Highlighting Tc25°C worms with 1hr hold at 15°C in pink (n=13 animals, 2 independent experiments); **(B”)** Highlighting Tc25°C worms with 4hr hold at 15°C in periwinkle (n=3 animals); **(B’’’)** Highlighting Tc15°C worms in blue (no additional temperature exposure) (n=12 animals). **(C)** For the highlighted data in **(B)**, AFD sensitivity to small temperature changes introduced by oscillations in the temperature stimulus (black), as measured by peak prominence. Significant calcium peaks labeled with ovals (see Methods section Quantification of AFD Sensitivity and Oscillation Prominence for more information). **(C’)** Peak prominence for each oscillation in the ramp in **(C)**, calculated as the difference (ΔR/R) in AFD soma calcium between the maximum and the preceding minimum. Although the temperature derivative experience is identical for each oscillation, note how the AFD responses to these oscillations vary across the stimulus in reference to the Tc.

We next asked whether this shifted response range reflects a discrete region for responsiveness or, instead, a graded gain landscape centered around an optimal temperature response region. To distinguish between these possibilities, we quantified peak prominence across the temperature range as a measure of response strength. Peak prominence represents the oscillation amplitudes to the temperature derivatives during the thermal ramps (Fig. 3C-C’). To more precisely characterize this relationship, we further refined this metric into a functional sensory gain, defined as the peak prominence normalized by the mean positive temperature derivative 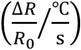 (Fig. S3E). By keeping the temperature derivatives constant but varying *when* the animals experience them relative to the cultivation temperature, we mapped AFD gain across the temperature space. We observed a ~4-fold modulation in the calcium response to the same temperature derivative, depending on the position of the derivative relative to the animal’s cultivation temperature preference (Fig. 3C-C’ and S3E). These measurements reveal that the AFD calcium response amplitude for a given temperature derivative dynamically changes across the temperature gradient relative to the absolute temperature preference. By scaling responses according to the position of the derivative relative to the animal’s preferred temperature, experience-dependent gain modulation ensures maximal sensitivity to thermal changes as the animal approaches its navigational goal.

To further examine this, we drew inspiration from Q10, a classical measure of temperature sensitivity (Běhrádek, 1930; Mundim et al., 2020). Q10 describes how strongly a process changes when temperature increases by 10°C, and has been previously measured for AFD by using physiologically-derived values or calcium-derived estimates under a single cultivation condition (Ramot et al., 2008a; Takeishi et al., 2016). Inspired by these studies, we designed experiments to investigate AFD experience-dependent response gain for animals raised at specific cultivation temperatures. In our experimental approach, we delivered three brief 0.5°C/min linear ramps (17-18°C, 21.5-22.5°C, and 25.5-26.5°C) for animals raised at 15°C, 20°C, or 25°C (Fig. 4A). These ranges were selected to induce robust responses for each cultivation temperature, considering the known lower bound of the response range for each condition. This experimental approach also enabled us to directly examine if the reduction of responses observed for the longer thermal ramps (Fig. 3) resulted from the eventual sensitization of the neuron to the duration of the thermal stimuli, or from a *bona fide* response range due to the experience-dependent gain modulation.

**Figure 4.**
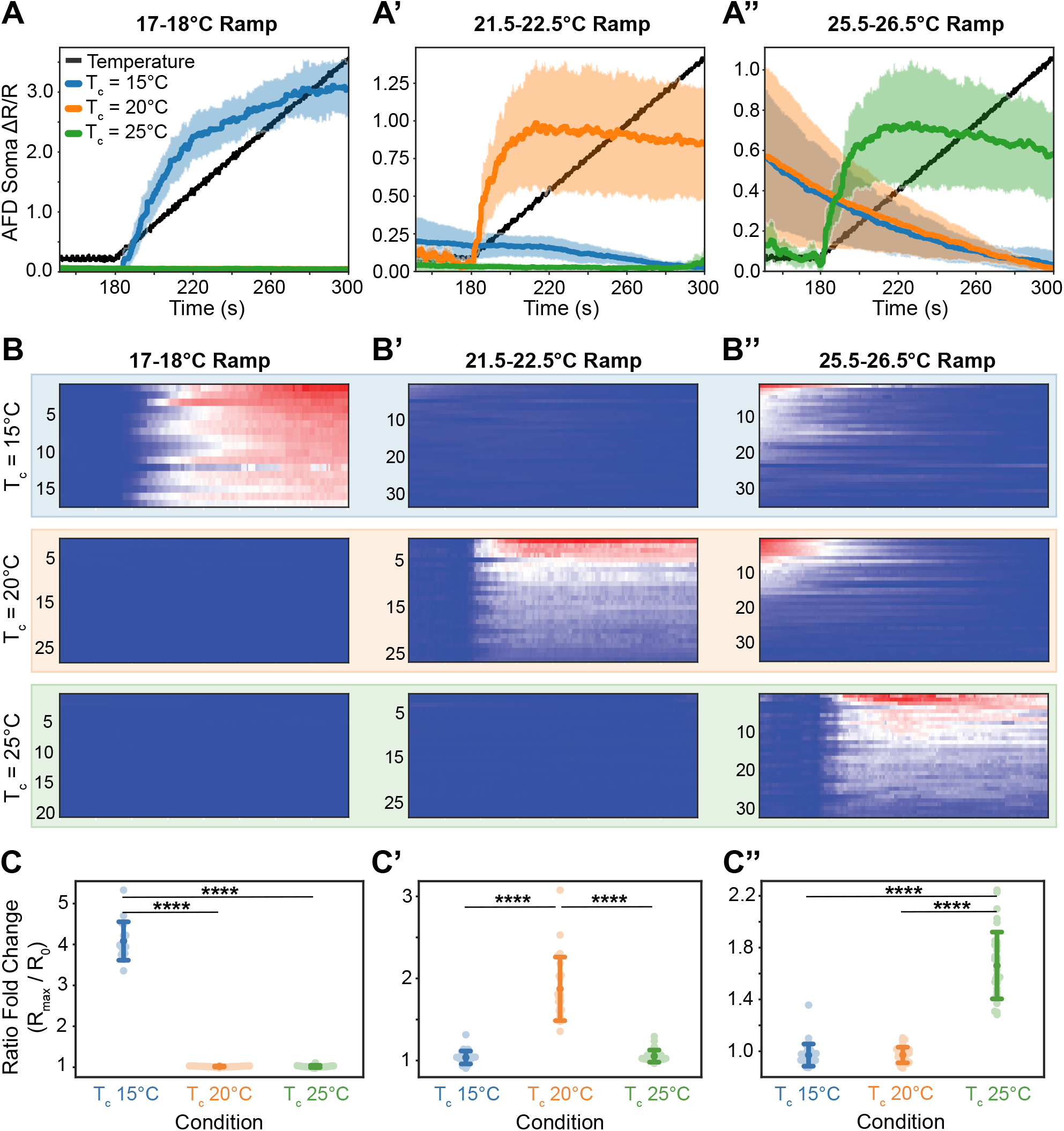
AFD exhibits increased calcium responses to the same temperature derivative around its cultivation temperature. **(A-A″)** Mean ratiometric AFD soma Ca^2+^ responses (ΔR/R) to short heating stimuli (delivered after a 3 minute hold at the starting temperature). Animals were exposed to a 2 minute upramp at 0.5°C/min (temperature stimuli shown with black line). Colored traces indicate animals trained at T_c_ = 15°C (blue), 20°C (orange), or 25°C (green). Darker line corresponds to mean; shading denotes ±1 standard deviation. **(A)** 17-18°C ramp (n=17 for Tc15;n=28 for Tc20 and n=20 for Tc25). **(A’)** 21.5-22.5°C ramp (n=33 for Tc15;n=26 for Tc20 and n=28 Tc25). **(A’’)** 25.5-26.5°C ramp (n=34 for Tc15;n=35 for Tc20 and n=32 for Tc25). As explained in the Methods section, a 3 minute holding time at the ramp’s starting temperature was included in all temperature protocols to stabilize the initial AFD response across conditions upon starting the experiment. Thus, the decay observed in (A’’) for Tc15 and Tc20 corresponds to the initial response to the temperature rise from Tc to 25.5°C, but no response to the ramp stimulus. **(B-B’’)**: AFD Ca^2+^ responses for single animals, shown as normalized heatmaps and corresponding to **(A-A″)**. Each row represents the response of a single animal over time. Blue indicates the minimum and red indicates the maximum calcium response observed for that experiment, for the indicated conditions. Note the response ranges of AFD based on the cultivation conditions and temperature preference of the animals. **(C-C″)** Quantification of the AFD responses to the upramps shown in **(A-A″)**. Points represent individual animals, and colored markers indicate the mean response for each condition (Tc15 in blue, Tc20 in orange, Tc25 in green) with corresponding error bars denoting ±1 standard deviation. Ramp-evoked responses are quantified as ratio fold change (R_max_/R_0_), where R_max_ is the maximum mean AFD response following ramp onset and R_0_ is the pre-ramp baseline (average ratio for the 30 seconds prior to the ramp). Statistical significance was assessed using a Kruskal-Wallis test followed by post hoc Dunn’s test;****denotes p <0.0001.

Sensitivity was measured via the fold change of the ratiometric signal, 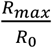, where *R*_*max*_ and *R*_0_ are the maximum ratiometric response to the stimulus and the baseline just before, respectively. Consistent with a sliding gain window, Tc25 animals exhibited a robust 1.66 ±0.26-fold increase in ratiometric signal in response to the 25.5°C ramp (Fig. 4A”, B’’,C’’), whereas Tc15 animals responded selectively to the 17°C ramp (4.08 ±0.47-fold increase) (Fig. 4A, B, C). Notably, neither Tc15 nor Tc25 animals responded strongly to the 21.5°C ramp (Fig. 4A’, B’, C’), indicating that responsiveness is not determined by absolute temperature alone but by the alignment of stimulus temperature with the gain window defined by cultivation temperature preference of the organism.

Together, these results indicate that AFD responses are governed by two temporally distinct processes: 1) a rapidly adapting sensory threshold operating on tens of seconds timescales and 2) a slower, experience-dependent gain modulation that shifts how temperature derivatives are represented by modulating the gain response in AFD based on where the derivative is experienced in the temperature gradient relative to the cultivation temperature goal. Moreover, our findings demonstrate how a derivative-based sensory code can represent absolute temperature through experience-dependent gain, allowing AFD to encode both temperature change and position relative to the preferred thermal setpoint. The combination of rapid leaky integration and slower gain modulation enables AFD to adapt across timescales while preserving sensitivity to the learned thermal preference.

To determine how these processes shape sensory encoding and behavior, we next developed a temperature playback paradigm that allows recent sensory experience to be dissociated from learned temperature preference and directly tests how long-term thermal history modulates AFD responses (Fig. 5A). In this setup, temperature trajectories recorded from freely moving animals were replayed to immobilized conspecifics. This approach provided a controlled, reproducible, and editable single-modality (temperature) virtual-reality experience faithfully derived from naturalistic behavior, enabling decoupling of sensory inputs from behavioral history. As such, it allowed us to probe AFD neural processing as if the animal were actively navigating while enabling precise experimental control to examine the influence of training conditions on AFD responses. To validate our approach, we first replayed naturalistic temperature trajectories to immobilized animals while recording AFD activity (Fig. 5A-B). Immobilized animals with the same training conditions as their freely moving counterparts exhibited indistinguishable AFD responses under comparable sensory contexts, as quantified by Pearsons’s correlation coefficient (0.96) and the mean square error (MSE, 0.014) between the responses. Our findings support previous observations that AFD faithfully encodes the temporal complexity of natural thermal experiences and achieves this independent of movement-related feedback (Tsukada et al., 2016).

**Figure 5.**
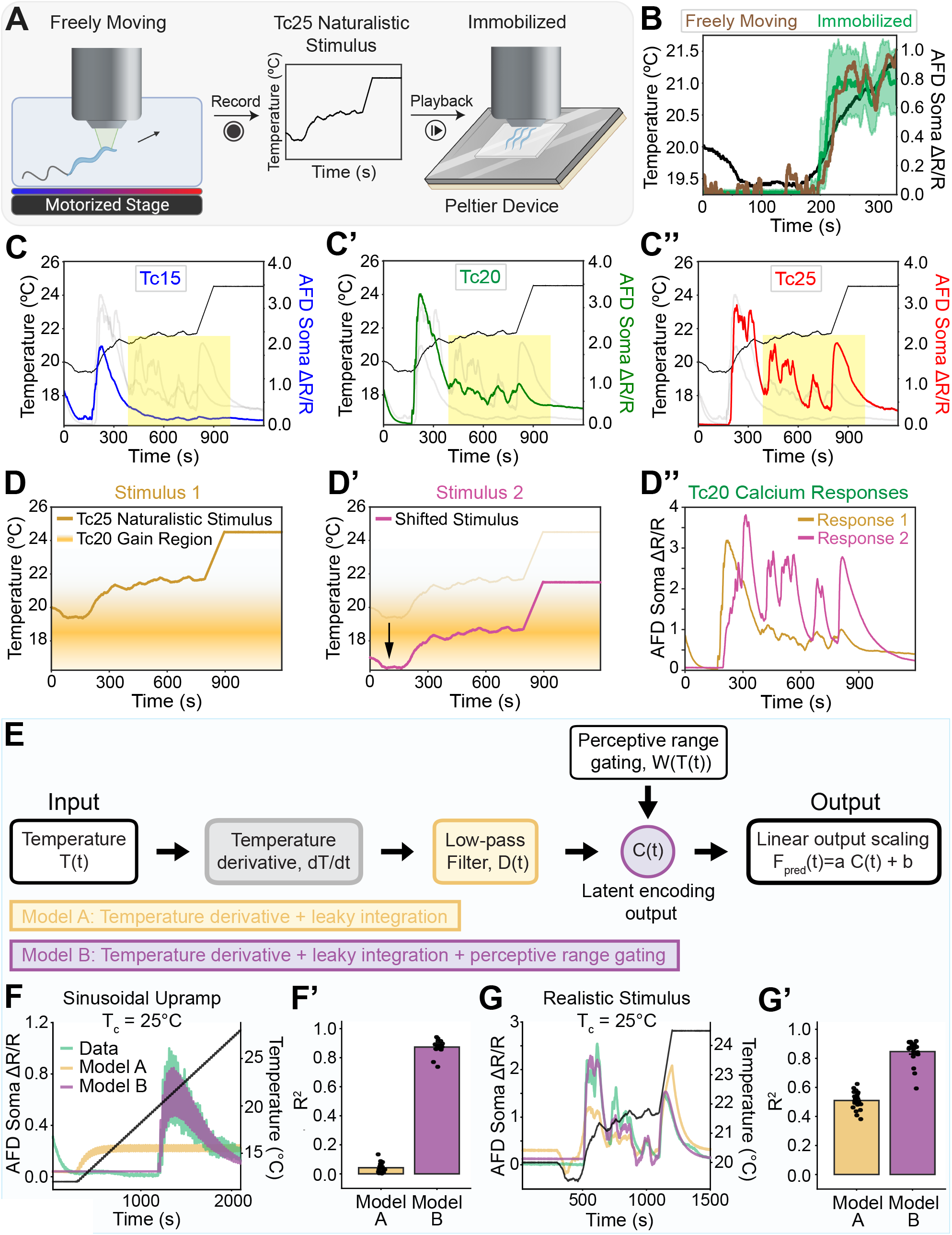
Replay of thermal experience reveals AFD’s context-dependent gain region modulated by temperature preference. **(A)** Schematic of the “temperature playback”protocol. A freely moving animal is tracked during a standard thermotaxis assay to obtain its x-y coordinates and resulting thermal experience. The identical temperature time course is then delivered on a fixed-temperature stage to an immobilized animal, enabling replay of a naturalistic stimulus while recording AFD Ca^2+^ (ΔR/R). **(B)** Comparison of AFD responses to the same temperature stimulus (black) in immobilized (green) and freely moving (brown) animals. **(C-C’’)** An identical stimulus is presented after 4h incubation at 15°C, 20°C, or 25°C. Each condition is highlighted in a different panel (Tc15 in **(C)** (n=13 animals, 3 independent experiments);Tc20 in **(C’)** (n=24 animals, 5 independent experiments);and Tc25 in **(C”)** (n=21 animals, 5 independent experiments)) with average responses recorded for AFD. Yellow box highlights the region in which we observed substantially different AFD responses based on cultivation temperature preference. **(D-D’)** Cartoon of the experimental design and results **(D”)** in which a naturalistic stimulus (based on a track of an animal trained at Tc25 and performing thermotaxis towards a warmer temperature, identical to that used in **(C-C’’)**) is replayed to an animal trained at Tc20 (like in **(C’)**). Briefly, in **(D)** the experience of the Tc25 animal (brown line, Stimulus 1) is displayed in the context of the predicted gain region for a Tc20 animal. The gold gradient bar denotes the temperature region of maximal AFD sensitivity, with the darkest shade marking the highest sensitivity. **(D’)** The same naturalistic stimulus is replayed to the Tc20 animals, but now the stimulus is down-shifted 3 degrees so that it falls in the predicted gain region (gain region depicted by gold band, and the experimental design called “Stimulus 2”). Temperature responses for AFD are recorded and compared in **(D”)**. Compare the response to Stimulus 2 (pink, called Response 2) with the response to Stimulus 1 (called Response 1, the same one shown in **(C’)**) and note the robust response to Stimulus 2 (n=27 animals, 3 independent experiments), when the preference-based gain region is aligned to the stimulus. **(E)** Schematic of a new mathematical model incorporating experience-dependent gain. After the low-pass filter step described in Fig. 2F, the latent variable is multiplied by the value of a gamma distribution as a function of temperature preference (Tc), then the final Ca^2+^ prediction is obtained by linear scaling. **(F)** For the thermal stimulus of Fig. 3 with Tc25-trained worms, the mean AFD soma Ca^2+^ (ΔR/R) (light green trace) is shown with the older model prediction (yellow, Model A) and new model prediction (purple, Model B)). **(F’)** For the experiment in **(F)**, individual fraction of variance explained for the AFD calcium trace of each worm, as predicted by Model A and Model B. **(G)** As in **(F)** for the thermal stimulus of **(C-C’’). (G’)** As in **(F’)**, for the experiment in **(G)**.

This paradigm enables an experimental design not possible in freely behaving worms. Unlike freely behaving worms, which inevitably execute their learned temperature preferences, this setup lets us train an animal to prefer one temperature and then immerse it in the naturalistic thermal experience of another animal that has navigated the gradient towards a different preferred temperature. We leveraged this feature, using this paradigm to dissociate the preference from the experience, thereby investigating how AFD encodes identical input streams in animals with different learned temperature preferences. Using thermal replay, we exposed animals with different training histories to the same precise temperature experience, enabling direct comparison of AFD encoding across conditions. To examine whether training conditions modulate AFD sensation, we exposed animals raised at Tc15, Tc20, and Tc25 to the same temperature trajectory derived from freely moving Tc25 animals (Fig. 5A). We observed distinct response strengths across groups. Specifically, Tc25 animals achieved the highest response strengths (Fig. 5C’’, yellow area), in contrast to Tc20 and Tc15 (Fig 5C’ and C, respectively). This indicates that AFD sensitivity decreased as the training temperature deviated more from the stimulus-corresponding experience (in this case, of Tc25). This effect was particularly evident in the highlighted region (yellow box, Fig. 5C-C’’), where the same thermal stimulus evoked different AFD responses depending on cultivation temperature preference.

We hypothesized that AFD gain control differentially represents temperature changes according to the temperature position relative to the cultivation temperature (Fig. 5D). If so, absolute temperature alone could not determine response magnitude; instead, sensitivity should depend on where a given temperature derivative falls within the gain window defined by the organism’s cultivation temperature preference. This prediction implies that for a fixed sequence of stimuli, the gain profile could be recapitulated by shifting the stimuli relative to the cultivation temperature preference. To test this, we designed an experiment in which we kept the cultivation temperature constant across conditions (Tc20), as well as the naturalistic temperature trajectory experienced by the animal (i.e., the experienced temperature derivatives). We then designed two trajectories for the thermal stimuli, the variable being the starting temperature of the stimuli, which in turn shifted the thermal position of the derivatives relative to the gain window for Tc20 animals (Fig. 5D, D’). We observed that taking a naturalistic track derived from Tc25 animals results in increased responses for Tc20 animals if those exact same stimuli were down-shifted 3°C and replayed to Tc20 animals (Fig. 5D’’). These results demonstrate that AFD gain control is a product of the relative position of the temperature derivatives to the cultivation temperature goal.

We next examined whether the linear stimulus-response mapping observed in freely moving animals near cultivation temperature (Fig. 2E) generalizes under these new conditions of varied temperature experience. We observed that although the Baseline Derivative Model accurately reproduced AFD responses under freely moving conditions near cultivation temperature (Fig. 2G), it performed poorly when tested on broader stimulus conditions. In particular, when animals were subjected to sinusoidal up-ramps or naturalistic stimuli across different cultivation temperatures (Tc15, Tc20, and Tc25), the Baseline Derivative Model provided poor fits (Model A in Fig. 5F,F’ and 5G,G’; R^2^=0.04 and R^2^=0.51, respectively). These low R^2^ values indicate that our previous model is insufficient to describe AFD responses across diverse temperature stimuli. We then examined the relationship between calcium and the filtered derivative drive across cultivation temperatures and stimuli and observed that the filtered derivative drive–calcium relationship did not follow a single linear mapping (Fig. S3F-G). Thus, while AFD responses in freely moving worms can be approximated by a linear model near cultivation temperature (Fig. 2E-H), this approximation does not generalize across broader thermal contexts and require non-linear relationships that account for the observed gain modulation of AFD.

To account for this, we developed a State-Dependent Gain Model (Mahéet al., 2015), which extends the derivative-based framework by incorporating a cultivation temperature-dependent gain function (Fig. 5E). As in the Baseline Derivative Model, the experienced temperature is first converted into its rate of change (dT/dt) and passed through a low-pass filter to generate a temporally integrated drive signal. This drive signal is then modulated by a temperature-dependent gain window, W(T(t)). We implement W(T(t)) as a gamma-shaped function that defines a temperature-dependent sensitivity profile. The gain peaks near the preferred temperature and decreases on either side. The parameters of W(T(t)), including its onset and shape, are inferred directly from calcium data. The resulting gated signal, *C*(*t*) = *W*[*T*(*t*)] ·*D*(*t*), is then linearly transformed to obtain the predicted calcium responses. We observed that the State-Dependent Gain Model substantially improved predictive accuracy across sinusoidal up-ramps and naturalistic stimuli conditions and outperformed the Baseline Derivative Model (Model B in Fig. 5F-F’ and 5G-G’; R^2^=0.88 and R^2^=0.78, respectively). Together, these results indicate that AFD calcium dynamics are best described by a combination of temporal processing of temperature derivatives and cultivation temperature-dependent gain modulation. Incorporating a gamma-shaped sensory window, defined as a unimodal, asymmetric gain profile with a gradual rise, peak sensitivity, and decay provides a compact and biologically interpretable mechanism for adaptive thermosensory encoding of behaviorally relevant sensory information across diverse stimulus regimes.

To understand the role of gain control in thermotactic behavior, we next examined behavioral strategies in the context of the observed AFD gain levels. We started by evaluating general migration of Tc25 animals performing thermophilic behavior. AFD’s gain modulation is highest at 22.5°C(Fig. 6A). We observed that, on average, Tc25 animals altered their locomotory strategies near 22°C(Fig. 6B-C), correlating with the AFD gain region for Tc25. We then quantified and compared locomotory strategies across the gradient by dividing our analyses of the gradient into two 2°C bins corresponding to 20-22°C(before the peak of the gain region) (Fig. 6F) and 22-24°C(containing the peak of the gain region) (Fig. 6G). We quantified the distributions of run angles at these defined temperature regions (Fig. 6F’ and 6G’) and observed that run angles are more oblique (closer to the 90°isothermal direction) at the temperatures closest to the thermal preference (compare Fig. 6F’ vs 6G’). This is consistent with a change of locomotory strategy, with animals moving perpendicular to the gradient. Our findings are consistent with previous studies showing that animals alter their locomotory strategies as they approach their preferred temperature, transitioning from gradient migration to isothermal tracking within approximately 2–3°C of their cultivation temperature (Chi et al., 2007; Garrity et al., 2010; Luo et al., 2006; Ryu and Samuel, 2002). We extend these findings by showing that AFD gain regions correlate with and might shape these distinct thermotactic behaviors.

**Figure 6.**
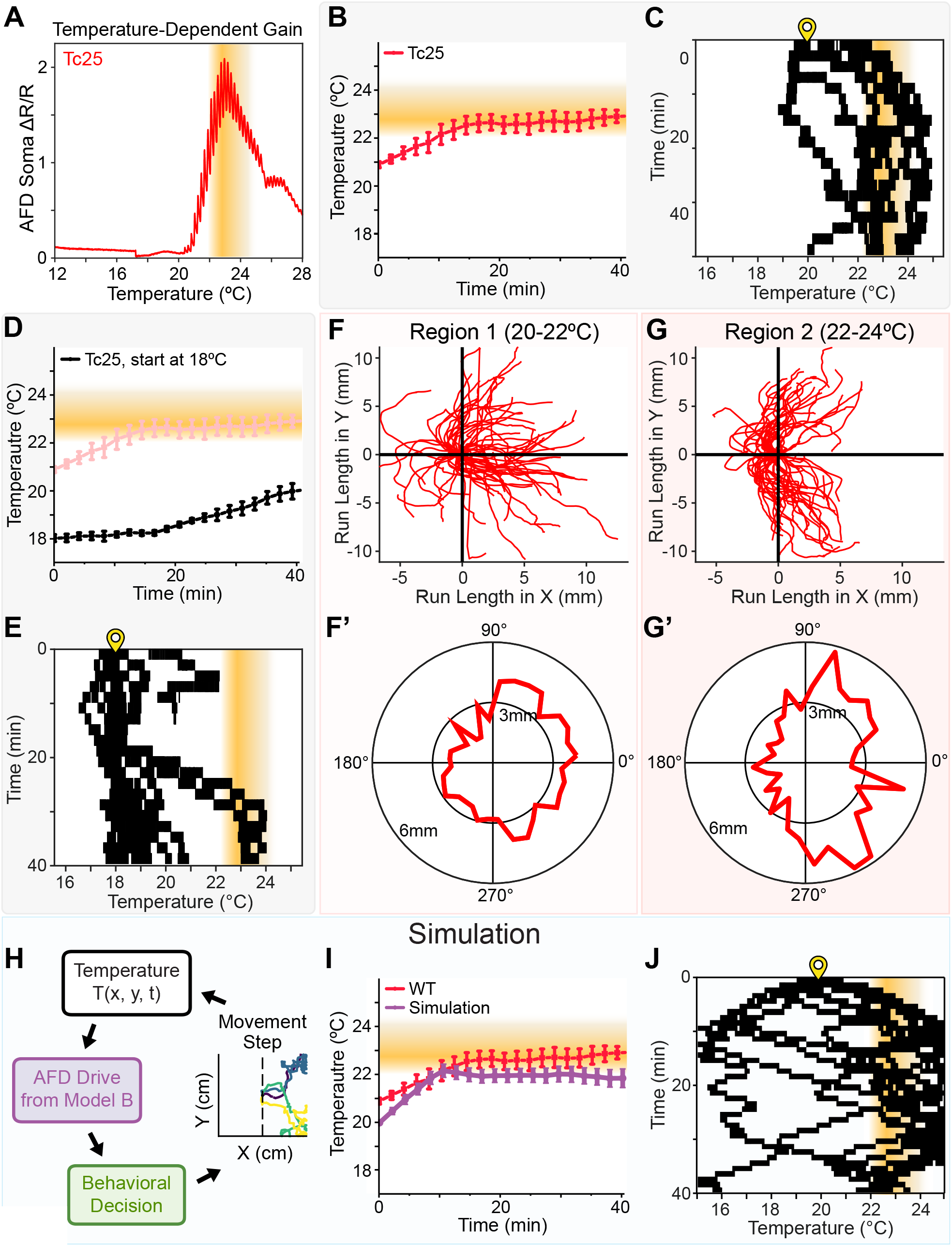
AFD’s temperature-dependent sensitivity predicts the gradient locations where animals transition from thermophilic migration to oblique movement and isothermal tracking. **(A)** Representative mean (n=9 animals) AFD soma Ca^2+^ response (ΔR/R) for Tc25-trained worms to the sinusoidal upramp thermal stimulus of Fig. 3 (dark line). Gold gradient bar denotes the temperature region of maximal AFD sensitivity, with the darkest shade marking the highest sensitivity. **(B)** Average thermotaxis movement of animals trained at Tc25 (n=45 animals, 5 independent experiments) placed on a linear 1°C/cm thermal gradient, starting at ~20°C. Gold gradient bar uses the temperature range from **(A)** to approximate the location of AFD’s maximal sensitivity on the gradient. Error bars denote SEM. **(C)** Raster plot of the position of the animals on the thermal gradient over time for the experiment in **(B)**. Yellow location pin marks the start temperature at which animals were initially placed. Gold gradient bar as in (A-B). **(D)** Data from **(B)** (pink) overlaid with behavior of Tc25 animals (n=36 animals, 4 independent experiments) that were placed at 18°C on an identical thermal gradient, and their average movement tracked (black line). Error bars denote SEM. **(E)** As in **(C)**, raster plot corresponding to the 18°C start condition in **(D). (F-G’)** Tracks of animals in two distinct temperature regions of the thermotaxis assay, corresponding to the gain region of AFD in Tc25-trained animals. Data are from eleven 15-minute thermotaxis assays on a 1°C/cm (15°C to 25°C along the horizontal X direction, where distance is proportional to temperature) gradient with Tc25 animals placed at 20°C. The 40 longest runs within each temperature bin were computationally identified and plotted with start points aligned to (0, 0). **(F)** Runs from the 20-22°C region. **(F’)** Polar histogram of mean run distance per 10°angular bin from the data in **(F)**. 0° represents forward movement directly toward higher temperatures, and deviations represent oblique movement. **(G)** As in **(F)**, for the 22-24°C region. **(G’)** As in **(F’)**, for the data in **(G). (H)** Simulation of thermotaxis behavior using the expanded AFD sensory model from Fig. 5. At each 1s time step, worm position on a simulated thermal gradient determines the temperature experience over time, which the AFD model uses to compute a latent drive variable that then maps to probabilistic behavioral decisions using parameters derived from wildtype behavioral assays (See Methods). Lower AFD drive promotes movement toward warmer temperatures, while higher AFD drive promotes isothermal-like movement (in this case, movement perpendicular to the gradient). The simulation was repeated for each worm at each time step to generate the tracks in **(I-J). (I)** Data from **(B)** (red line) overlaid with simulated behavior of animals (purple line; n=500 worm simulations) starting at 20°C. Red error bars denote SEM, and purple bars denote 95%confidence intervals. **(J)** As in **(C)**, showing 15 randomly selected simulated worms from the data in **(I)**.

We further examined the relationship between the gain regions and the locomotory strategies by localizing animals outside the gain areas of AFD (at 18°Crather than 20°C) and examining thermotactic behavior (Fig. 6D). We observed that Tc25 animals that started at 18°Cwere defective in their capacity to move towards the warmer temperatures until they entered the gain region of AFD (Fig. 6D-E). These observations further support a model in which experience-dependent increases in AFD gain could contribute to the behavioral strategies of the organism, extending prior work showing that AFD activity and output are sufficient to shape distinct thermotactic behaviors (Kuhara et al., 2011; Luo et al., 2014).

Lastly, we asked whether the AFD encoding model, when coupled to simple sensorimotor rules, was sufficient to reproduce key features of thermotactic behavior. To do this, we built a minimal behavioral simulator that linked the sensory gain framework to strategy selection probabilities (Fig. 6H). In the simulator, the instantaneous temperature derivative experienced by a simulated worm was passed through Model B (Fig. 5E), and the resulting output was scaled by sensory gain to generate a behavioral drive (Fig 6H). The magnitude of this drive determined the probability of different movement strategies, with stronger drive values biasing the animal to move perpendicular to the gradient rather than continued migration along the gradient. Iterative simulations produced tracks, which were compared to the empirical data of real Tc25 animals navigating across the gradient (Fig. 6I–J). The simulation reproduced key aspects of the behavioral switch observed in our experimental data (Fig. 6I). Specifically, the simulation captured three qualitative features of the experimental data: an initial warmward migration, a change in locomotory strategy as animals approached the sensory gain region near ~22°C, and an enrichment of simulated trajectories running obliquely to the gradient near this temperature range. This suggests that a model combining derivative encoding, experience-dependent gain, and simple strategy-selection rules can capture core features of thermotactic navigation.

## Discussion

In this study we find that responses of the primary *C. elegans* thermosensory neuron AFD are governed by two interacting processes: a seconds-scale temporal integration process that tracks recent temperature changes, and hours-scale experience-dependent gain modulation that shifts the range over which temperature changes are strongly represented in AFD calcium responses. Prior studies have documented several of these striking, and at first glance paradoxical, properties of AFD, including derivative-like responses, rapid adaptation, and experience-dependent shifts in operating range (Clark et al., 2007b, 2006; Kimura et al., 2004; Ramot et al., 2008a; Tsukada et al., 2016; Wasserman et al., 2011; Yu et al., 2014). The recurrence of these features across studies using different stimuli, recording methods and analytical approaches underscores their robustness. At the same time, integrating these observations into a unified encoding framework has remained challenging, in part because AFD appears to encode different features of the thermal experience on different timescales. For example, the term “operating range”has often been used to describe AFD’s cultivation-dependent response window, encompassing both the threshold for activation and the temperature range over which the neuron responds (Kobayashi et al., 2016; Wasserman et al., 2011; Yu et al., 2014). Our contribution is to show that these properties can be decomposed into separable computations, like temporal integration, threshold adaptation, and gain modulation, that together explain how derivative signals become goal-relevant. By combining calcium imaging in freely moving animals performing thermotaxis with a replay paradigm using naturalistic thermal trajectories, we separated these contributions into components with distinct timescales and functional roles: a rapidly shifting response threshold that tracks recent thermal history, and a slower, experience-dependent gain function that biases sensitivity toward temperatures near the animal’s learned cultivation temperature. We then formalized these features in a unified mathematical model and tested how they account for AFD responses across stimulus regimes. Together, our results show how AFD can remain fundamentally derivative-sensitive while supporting behavior anchored to an absolute, learned temperature preference: seconds-scale temporal integration preserves sensitivity to recent temperature changes, while slower cultivation-temperature-dependent gain weights those derivative signals according to their relevance to the animal’s learned thermotactic goal.

Foundational concepts such as efficient coding, linear–nonlinear receptive-field models, and normalization or gain-control frameworks have shaped sensory neuroscience by providing compact explanations for how neural systems extract features, reduce redundancy, and maintain sensitivity across changing stimulus statistics (Barlow, 1961; Carandini and Heeger, 2011; Chichilnisky, 2001; Schwartz et al., 2006; Simoncelli and Olshausen, 2001). Prior modeling in *C. elegans* has similarly shown how simple models can identify interpretable internal variables linking sensory history and neuronal responses, including in AFD thermosensation (Batra and Sharma, 2026; Clark et al., 2007a; Mobille et al., 2023; Tsukada et al., 2016). Related models in other *C. elegans* sensory systems have further shown how such internal variables can predict navigation and behavioral strategy selection (Gray et al., 2005; Kato et al., 2014; Levy and Bargmann, 2020). Viewed alongside these studies, our findings suggest that sensory neurons may reuse related computational motifs across modalities: temporal filtering to extract stimulus dynamics, adaptive variables to encode recent experience, and linear-nonlinear (LN) transformations that regulate the influence of sensory signals on navigation. LN models in particular are a useful framework in sensory neuroscience for relating stimulus features to neural responses (Chichilnisky, 2001; Schwartz et al., 2006). Consistent with the linear transformation in this framework, we find that AFD responses within a narrow thermal range near cultivation temperature are well approximated by a linear description of derivative-driven activity. However, this local approximation does not generalize across broader thermal contexts. Instead, we find that the relationship between filtered derivative drive and calcium response is not single-valued, indicating that no single static nonlinearity can account for the observed responses. AFD responses are better described by a state-dependent model in which a cultivation-dependent gain function rescales responses to temperature derivatives according to their position relative to the learned temperature goal. The molecular mechanisms that implement the gain function remain to be determined, but recent work on AFD thermotransduction and rapid adaptation provides candidate pathways through cGMP, calcium feedback, receptor guanylyl cyclases, and experience-dependent regulation of thermosensory machinery (Harris et al., 2026; Hill and Sengupta, 2024; Inada et al., 2006; Takeishi et al., 2016; Wang et al., 2013; Yu et al., 2014). Together, our findings suggest that AFD gain control resembles canonical sensory gain-control mechanisms in rescaling sensitivity but differs in that the gain function is referenced to a learned behavioral setpoint. This architecture allows a derivative-based sensory code to support navigation toward an absolute temperature goal and may represent a broader sensory strategy for aligning dynamic sensory encoding with learned behavioral targets.

In the broader sensory-coding landscape, gain control is often understood as an adaptive rescaling of neural sensitivity that matches responses to stimulus statistics and biophysical constraints, thereby preserving dynamic range under variable conditions. Classical examples include contrast gain control and normalization in vision, divisive normalization in olfaction, and sound-level or contrast adaptation in audition (Carandini and Heeger, 2011; Laughlin, 1989; Olsen et al., 2010; Rabinowitz et al., 2011). In *C. elegans*, sensory responsiveness is also shaped by circuit state and experience, including calcineurin-dependent sensory gain control, network-state feedback onto olfactory responses, and normalization-like computations in sensory choice circuits (Cohen et al., 2019; Gordus et al., 2015; Ikejiri et al., 2023; Kuhara et al., 2002). In contrast to these canonical formulations, the gain modulation we observe in AFD is a *setpoint-referenced gain control*: the high-gain regime shifts with cultivation temperature, preserving the same functional relationship across different absolute temperatures and effectively aligning derivative sensitivity with error, and relative to the learned goal. This coding scheme amplifies derivative-evoked responses near the learned setpoint, and attenuates them outside it, over-representing the thermal changes most relevant as the animals approach the preferred temperature range. Thus, AFD represents a new example of sensory gain control with a gain anchored around a learned setpoint, rather than other canonical sensory gain controls that are scaled around global stimulus intensity or stimulus statistics.

Our results also support a model in which AFD’s gain-weighted temperature-change signals influence locomotory strategy during thermotaxis behavior. During thermotaxis, animals migrate across a gradient to approach their preferred cultivation temperature and then transition to isothermal tracking near that goal (Hedgecock and Russell, 1975; Iino and Yoshida, 2009). AFD’s major postsynaptic partner is AIY (White et al., 1986), and the AFD–AIY synapse transmits graded excitatory signals (Narayan et al., 2011). AIY activity has been implicated in promoting forward locomotion and reducing reorientations during navigation (Kocabas et al., 2012). In this circuit context, our model suggests that AFD could provide a context-dependent drive onto AIY that tunes the balance between turns and runs as animals move through different regions of the gradient. As animals approach the cultivation temperature and enter AFD’s high-gain regime, we observe a shift toward trajectories more perpendicular to the gradient. We hypothesize that, in this regime, even shallower temperature derivatives generated during oblique or isothermal-like runs can produce sufficient AFD activation to recruit AIY and promote persistent forward movement. Outside the gain region, the same derivatives would evoke weaker AFD responses and may be less effective at driving this transition. Thus, experience-dependent AFD gain could provide a mechanism for switching from gradient migration to isothermal tracking near the learned temperature goal. This interpretation is supported by a minimal behavioral simulator that links the sensory gain framework to strategy-selection probabilities. The simulator does not capture the full downstream thermotaxis circuit, but it tests whether the sensory transformation we identify is sufficient, when coupled to simple locomotory rules, to reproduce key behavioral transitions. The fact that this minimal simulation reproduced key aspects of the behavior observed in our experimental data supports a model in which derivative encoding, experience-dependent gain, and simple strategy-selection rules capture core features of thermotactic navigation.

This framework also provides a way to interpret genetic perturbations that alter thermotactic strategy. For example, in *inx-1* mutants, AIY becomes hyper-responsive because circuit filtering is reduced. *inx-1* mutants also promote isothermal tracking outside the cultivation temperature window (Almoril-Porras et al., 2025). We propose that wild-type gain modulation and *inx-1*-dependent AIY hyper-responsiveness may converge on a common functional variable: the promotion of runs oblique to the gradient due to the higher probability of AIY activation. In wild-type animals, AIY may be preferentially recruited near the goal because AFD’s setpoint-referenced gain amplifies derivative-evoked input in the Tc-aligned regime. In *inx-1* mutants, AIY becomes hyper-responsive through altered circuit filtering, responding to smaller signals from AFD, even outside the gain region. In both cases, the probability of activating AIY in the context of a given temperature derivative influences run probability, including run probability at oblique angles to the gradient. These oblique angles to the gradient would result in smaller temperature derivatives when they are outside of the gain region (in wild type animals). But when in the gain region, or due to AIY-hyper-responsiveness (in *inx-1* mutants) they would elicit a response and therefore, run persistence in the isothermal strategy. Thus, gain modulation at either the sensory stage (AFD) or interneuron stage (AIY) may reshape thermotactic strategy by reweighting how thermal change signals are translated into turn/run decisions to a given temperature derivative.

More broadly, our results show how a sensory neuron can use gain control to embed an absolute behavioral goal into a derivative-based code. Many sensory systems preferentially encode changes, contrasts, or temporal derivatives, yet behavior often requires reference to stable goals or setpoints. AFD’s coding scheme allows a single sensory neuron class to extract multiple features from the same stimulus stream, such as recent change, local trajectory, and distance from goal, and convert them into navigation-relevant signals. Such setpoint-referenced gain control may represent a general strategy by which sensory neurons align dynamic encoding with learned behavioral goals.

## Acknowledgements

We thank Thierry Emonet for his thoughtful comments on this project and members of the Colón-Ramos lab for their feedback. The N2 strain was provided by the CGC, which is funded by NIH Office of Research Infrastructure Programs (P40 OD010440). Some of the diagrams used in the figures are modified versions of BioRender. This work was partially conducted at the Marine Biological Laboratories at Woods Hole, under a Whitman Research award to D.A.C.-R and H.S, and was inspired by work initiated during the Neural Systems and Behavior course. We thank the students and instructors from that course for fruitful discussions and inspiration, in particular Ifedayo-Emmanuel Adeyefa-Olasupo. Support to J.B. was provided by the Shurl and Kay Curci Foundation Award of the Life Sciences Research Foundation (LSRF). Support for J.D.H. was provided by the Ruth L. Kirschstein NRSA (NIH-NIMH-F32MH105063). This work was supported by the Howard Hughes Medical Institute (to A.L. and H.S.), including a Gilliam fellowship to E.C.-B. (GT11399) and a Hannah Gray Fellowship to A.C.C. (GT15993). The work was also supported by the National Institutes of Health grants to D.C.-R. (R35NS132156 and R01NS076558) and D.C. (NIH R01EY026555).

## Resource Availability

### Lead Contact

Further information and requests for resources and reagents should be directed to and will be fulfilled by the lead contact, Daniel Colón-Ramos.

### Materials availability

Plasmids and strains used in this study are available upon request to the lead contact.

### Data and Code Availability

- All behavior raw data as csv files can be found on GitHub including the corresponding analysis codes:
  - Deep Learning Repository: https://github.com/colonramoslab/CabezasBou-et-al.-2022.git
  - Repository with the codes for extracting animal coordinates from standard thermotaxis assays: https://github.com/colonramoslab/Almoril-Porras-et-al.-2023-Inx-1-.git;DOI: https://doi.org/10.5281/zenodo.14170612
  - All other analysis codes and data used in the article: https://github.com/colonramoslab/Diaz-Garcia-and-Beagan-et-al.-2026-AFD-.git
- Freely moving recordings and immobilized calcium-imaging acquisitions are available on Yale Dataverse: https://doi.org/10.60600/YU/H7BQYW
- Any additional information needed to reanalyze the data reported here can be obtained from the lead contact.

## Materials and Methods

### Temperature Training for All Assays

The day before each experiment, larval stage 4 (L4) animals were cultured with food overnight at 20°C. The next day and 4hrs before the experiment, the plates were shifted to an incubator at the corresponding training temperature, along with a plate without food and M9 buffer which were used later to transfer the animals into the behavioral arena for thermotaxis assays. For calcium imaging, a similar procedure was used with an imaging pad for recording of calcium dynamics in immobilized animals. For the latter, an aliquot with Levamisole was also included in the incubation with the objective of equilibrating all reagents the animals are exposed to, at the training temperature.

### Temperature Training for Up and Down Shift Experiments

For the specific case of the temperature downshift experiments, animals were cultivated at 20°C, then trained for 4hrs at 25°C. After this initial 4hr temperature training, animals were shifted to 15°Cfor different amounts of time: 0mins (referred to as Tc25°C), 1hr and 4hrs. A separate control group in this experiment followed the standard protocol of 20°Ccultivation and then trained for 4hrs at 15°C(referred to as Tc15°C). For all conditions, animals were then mounted for Calcium imaging as described in the Methods section: Imaging of Calcium Dynamics in Immobilized Animals. For the temperature upshift experiments, animals were cultivated at 20°C, then trained for 4hrs at 15°C. Following this initial 4hr temperature training, animals were shifted to 25°Cfor different timeframes as indicated in the results, including 5mins, 20mins and 1hr.

### Temperature Controlled Closed Loop System For Freely Moving Animal, Tracking Position and Calcium Dynamics Simultaneously

The temperature-controlled stage which generated a linear thermal gradient was custom built and adapted to fit under an upright Leica DM6 M Epifluorescence microscope with an X,Y motorized stage and a motorized nosepiece Z drive. The temperature gradient was generated similar to the previously reported steep gradient system (Almoril-Porras et al., 2025), where two thermoelectric Peltier modules that were independently set to the target temperatures via a proportional-integral-derivative (PID) controller (FTC200, Accuthermo) were used. This system in contrast was miniaturized to fit under the microscope, having each Peltier module mounted under one end of a 10cm x 3cm x 0.5cm metal plate with a thermal probe on top (SRTD-2, Omega) for feedback control. Excess heat was constantly removed using the same custom-built cooling system also reported previously (Almoril-Porras et al., 2025; Hawk et al., 2018). Once a stable linear gradient was set, an agar pad was cut (7cm x 3cm), placed over the metal plate, and allowed to reach the desired temperature (usually taking less than 10min). The precise gradient was confirmed by measuring the temperature across the agar pad, first with a separate T-type thermal probe (IT-24P, Physitemp) connected to a hand-held thermometer (HH66U, Omega), and then with an infrared temperature probe (Actron).

A single worm, trained at a specific temperature, was then transferred onto the agar pad at 20°C (which corresponds to the center of the gradient) using a micropipette in 3µL of the standard buffer solution M9, which was also prewarmed at 20°C. Immediately after crawling out of the M9 droplet, the worm was manually set in focus and centered within the field of view (FOV).

Fluorescence time-lapse imaging with 1ms exposure through a 5X/0.15NA FLUOTAR HCX PL APO air objective was automated via the Micro-Manager software. AFD neuron co-expressed GCaMP6 (green channel) and TagRFP (red channel) fluorescent proteins, and a Mightex LED Control Module (BSL-SA) was used to image the worms. Dual-channel recordings at 33FPS were achieved by incorporating a W-View Gemini image splitter (Hamamatsu, A12801-01), which simultaneously projected green and red fluorescence images onto the two halves of the Hamamatsu ORCA-Flash4.0 LT C11440-42U camera mounted on the microscope. The camera features a 33FPS framerate and pixel size of 6.5µm. When coupled with the 5x Plan APO objective, this configuration yielded an effective pixel size of 1.3µm at the sample plane. Based on the Nyquist principle, the practical spatial resolution of this imaging system is ~2.6 µm. The xy position of the motorized stage was recorded in the metadata of each image. To obtain real-time recordings for over tens of minutes with high temporal resolution, while at the same time avoiding CPU RAM overload, images were saved in OME-TIFF stack files directly to a solid-state drive (SSD).

To continuously image animals performing thermotaxis behavior, acquisition was initiated along with an automated, custom-built neuron tracking system coded in Beanshell for moving the stage while tracking the animals (See the Data and Code Availability section for repository link). The script tracked the xy movement of the AFD neuron by monitoring a bright spot in the neuronal TagRFP image with a consistent intensity. Every 500ms, the tracker checked the neuron’s xy coordinates, paused the acquisition, and quickly updated the xy stage’s position to recenter the neuron in the FOV before recording resumed. In this way, the script was able to keep the worms within the FOV while avoiding motion-blur artifacts. Because the stage’s xy position was recorded for each image, both the global movement of the worms, as well as the exact temperature they experienced could be deduced in post-analyses. In addition, the script ensured the neuron stayed in focus by deducing the optimal focus based on the current and two extra images acquired above and below the current focal position every 500ms. This automated 3D neuron tracker allowed us to record the calcium activity of the freely moving worms performing thermotaxis using a single camera setup with the requisite spatiotemporal resolution (2.6µm at 33Hz) necessary for tracking thermotaxis behavior and calcium dynamics in single neurons of freely moving animals until the animals reached their preferred temperature (which took approximately 10-20min). This is a significant increase in duration and temporal resolution of recordings when compared to the 2min at 5Hz manual tracking recordings by (Clark et al., 2007a). Furthermore, our system provides an improved spatial resolution compared to the 30 Hz dual-camera setup (0.5 NA, 2X objective) lacking z-axis auto-focus described by (Tsukada et al., 2016), which was digitally under sampled and limited to a practical spatial resolution of 16.0 µm., affording us larger quantitative datasets for analyses.

### Deep Learning Image Processing of Freely Moving Animal Thermotaxis Data

The large datasets necessitated automated methods that enabled data extraction and analyses of both the temperature experience, the worm locomotory strategy and the calcium dynamics in single neurons. Recorded videos were processed and analyzed using a custom deep neural network system (See the Data and Code Availability section for repository link). First, raw dual-channel video files (.ome stacks) for a single worm (SupFig1) were build pixel-to-pixel-registered dual-channel images from adjacently co-captured camera subarrays and visualized in full using a custom macro code in CytoShow (ImageJ) (Duncan et al., 2019). Videos where the neuron exited the field of view were identified and discarded manually, whereas out-of-focus videos were flagged using a machine learning algorithm. Specifically, the model was trained to detect only in-focus neurons utilizing the red fluorescent channel. Out-of-focus frames exhibited a characteristic drop in RFP signal, causing the model to register a loss of detection and automatically exclude the corresponding frames. If the number of frames without a neuron exceeded 10%of the total number of frames within the video, then the video was discarded. Out of 80 videos, 11 were manually discarded while 7 videos were automatically discarded by the model, underscoring the robustness of the tracking system and prediction model. Remaining videos were then processed by first overlaying green and red channels with pixel precision, and the accounted offset was used to split and save the raw video as two single channel image time-series.

We built and trained a U-Net, fully convolutional neural network (FCN). Our U-Net classified each pixel as either part of the neuron’s soma or dendrite, or as a background pixel. The probabilities across categories per pixel summed to one by design, and the highest probability category was chosen for that pixel. with the highest value indicating the most signal for that particular category. Our U-Net inputs one image with two color channels: the green and red fluorescent channels depicting the soma and dendrite of the AFD neuron, respectively (SFig1). Both channels were used to train the U-Net, which outputs a probability map over three categories (background, soma, and dendrite) for each pixel. Our CNN followed a traditional U-Net architecture. The model contained 27 layers in total: 18 convolution layers, 4 down-sampling layers, 4 up-sampling layers, and a concluding softmax layer to assign probability values for each of the three categories per pixel, totaling 309,075 tunable parameters (SFig1).

We trained our CNN using a small set of images (<300 frames) that were manually annotated to distinguish subcellular compartments of the AFD neuron in both green and red fluorescent channels. Model parameters were tuned by using 3 images per frame in the set: the green fluorescent channel, red fluorescent channel and the manually annotated mask where both pixels pertaining to the neuron dendrite and soma were marked. The loss function guided the tuning of parameters during model training; we used a form of the *F*_*β*_ function as a basis for our loss function. The *F*_*β*_ function traded off prioritizing precision (i.e. penalizing false positives) or recall (i.e. penalizing false negatives) via the *β* parameter.

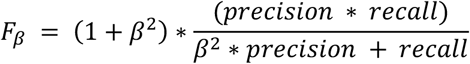

We set *β* = 2 to bias the model towards prioritizing recall, as we required a continuous segmentation of the dendrite, which often appeared dim along portions of the neuron. The loss function was then a weighted sum of *F*_2_ losses across the three categories as written below, where *w*_*j*_ are weights on each category, *y*_*j*_ are the ground-truth pixels corresponding to class *j*, and 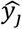 are the model predictions for class *j*. We used weights *w*_0_ = 1, *w*_1_ = 2, *w*_2_ = 30 to further prioritize dendrite detection. The model was trained using the Adam optimizer with a learning rate of 2.5 ·10^−4^.

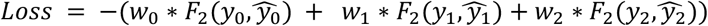

Our CNN used individual green and red fluorescent channels as input to predict a binary AFD skeleton mask for every image (SFig.1), which was then used to precisely quantify the location and pixel intensity of the GCaMP and RFP in all parts of the neuron. Since the model was trained to detect AFD’s soma as separate from the dendrite structure, our analysis was able to quantify neuronal activity with subcellular resolution.

### Temperature Controlled Closed Loop System for Tracking of Freely Moving Animals during Standard Thermotaxis Assays

The same thermotaxis behavior assay system previously reported was used (Almoril-Porras et al., 2025). It consists of 2 pairs of thermoelectric Peltier plates under an aluminum slab, one pair at the left and another at the right edge. They are controlled by two H Bridge amplifiers (Accuthermo FTX700D) in turn modulated by 2 PID controllers (Accuthermo FTC100D). The plates are directly cooled by a closed loop system circulating refrigerant cooled by a radiator in contact with dry ice. At each end a thermistor (Accuthermo TR2252) is fixed to the upper surface of the metal slab, approximately 0.25 inches into the gradient with respect to the Peltier modules. Each thermistor is connected to a PID controller. A square array of red LEDs surrounding the arena create a high contrast view of the animals, seen with bright outlines. A MightEx camera (BCE-B050-U) was used to capture images at 2FPS for 60mins. This entire system is enclosed in a modified cabinet. The main aluminum slab over which the thermal gradient is established is 10cmx10cm.

### Thermotaxis Behavioral Assay

Thermotaxis behavioral assays (without calcium imaging of AFD) were performed as previously described (Almoril-Porras et al., 2025; Hawk et al., 2018). An Agar pad, cut to a 10cmx10cm square, was directly placed over the metal slab surface and allowed to reach the desired temperatures for 5-10mins. Temperatures were independently validated before the experiment by using a high accuracy thermometer (Fluke 54-2B) with their standard thermocouple probes (80PK-1 bead). Just before experiments, animals were taken out of the incubator and gently picked into the plates without food, allowed to move briefly and then the plates were flooded with M9 buffer. Animals were then transferred in 3-4 M9 buffer droplets pipetted unto the midline of the pad (unless otherwise noted, this corresponds to the 20°C isothermal region), where each droplet carried 3-4 animals each. Animals were monitored via a top-mount digital camera MightEx (BCE-B050-U) until most of them exited from the droplet area. Immediately after, acquisition at 2FPS was started for the duration of the assay (60mins) as animals performed thermotaxis.

### Closed Loop System for Imaging Calcium Dynamics in Immobilized Animals

As previously described (Almoril-Porras et al., 2025; Hawk et al., 2018), a cooling system removing excess heat runs through a copper cooling block attached to the bottom surface of each Peltier plate. A custom LabView (National Instruments) software, H-bridge amplifier (Accuthermo FTXD700D) and controller (Accuthermo FTC200) that obtain temperature measurements from a thermal probe fixed on the plate (Omega SRTD-2) were used to control temperature of the stage. A Leica DM5500 and DM6B microscope with a 10X/0.40NA HC PL APO air objective, Photometrics Dual-View 2 (DV2) optical splitter, and a Hamamatsu ORCA-Flash 4.0 LT C11440-42U camera were used to simultaneously acquire images on both green and red emission channels for ratiometric signal analysis. MicroManager was used to acquire fluorescence images over time at 250ms exposure time, 1FPS, 4×4 binning, and with Z refocusing every 60s (Edelstein et al., 2014). Under this optical configuration, the 4×4 binning yielded an effective image pixel size of 2.6 µm, resulting in a practical spatial resolution of 5.2 µm at the sample plane. BioLED Light Source Control Module 470nm and 560nm powers were set to 3%and 10%, respectively. The temperature protocols were loaded into the custom LabView interface from .txt files containing the coordinates at 0.1s intervals.

### Imaging of Calcium Dynamics in Immobilized Animals

As previously described (Almoril-Porras et al., 2025), a large drop of warm 5%Agarose dissolved in M9 buffer was placed on a 22mmx22mm glass slide and pressed with a larger 25cmx75cm microscope slide to create the imaging pad. 18*μL* of Levamisole were placed over the pad. Worms were taken out of the incubator along with the reagents and picked from their respective cultivation plate unto one without food, removing excess bacteria afterwards manually and by letting the animals move briefly. After flooding the plate with M9, as many animals as possible were pipetted unto the pad in 2 droplets of 3 μL each. To enrich for dorsoventral view, a 5mins rest time was given before proceeding. Any excess liquid remaining was removed by slightly tilting the coverslip while keeping the worms uphill and then removing the excess on the bottom by cutting that section, pipetting the liquid out or lightly touching a corner with a wipe. The worms were grouped together using an eyelash pick. Once the pad was almost dry, the top coverslip was carefully placed at an angle over the pad. The imaging pad was then moved to the temperature stage, which was previously set to hold at the corresponding cultivation temperature while the image was then focused to start acquisition (~5mins). The holding temperature was validated before placing the imaging pad with the same high accuracy thermometer used for the standard thermotaxis assays but probing with a surface thermocouple (Omega SA3-T-SRTC). Acquisition was initiated by first starting image collection at 1FPS in MicroManager, and immediately after starting the protocol in the LabView script interactive window. At the end of each experiment, the LabView script returns a text file with the temperature readings every 0.1s and MicroManager a time stack in .tif format. Importantly, for stimuli such as the sinusoidal upramp (Fig. 3) and brief linear ramps (Fig. 4), a 3 minute and 5 minute hold at the initial temperature of the ramp was included in the protocol. This allowed AFD responses to stabilize after experiencing a shift from the pre-set hold at Tc while the imaging conditions were entered and the initial temperature of the ramp was established.

### Quantification and Statistical Analysis

#### Quantifying Neuronal Activity from Freely Moving Worms via Calcium Imaging

Quantification of calcium activity at the neuron’s soma was obtained by taking its average intensity from the GCaMP6s (Chen et al., 2013) green fluorescence and comparing it to the calcium-independent signal of the RFP. This ratiometric normalization was computed by first taking the ratio between GCaMP and RFP fluorescence signal at each timepoint R_t_ and then computing its rate of change by obtaining its first derivative (ΔR/R = (R_t_ - R_t-1_)/R_t-1_).

#### Quantifying Behavior in Freely Moving Animals Simultaneously Recorded for AFD Activity

Reversals, positive and negative runs, omega turns and pauses with local searches were manually annotated by observing animal movement over the acquired videos. The frame numbers were annotated, from which the exact time in the experiment and coordinates corresponding to these behaviors were obtained using a custom Python script (see the Data and Code Availability section for the repository link). AFD calcium plateaus were annotated based on oscillations sustained for 2 or more periods around the local baseline. The temperature derivative experienced during plateau events was calculated from: 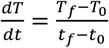, where the subscripts f and 0 denote the final and start of the plateau, respectively. The plateau duration was defined as *t*_*f*_ −*t*_0_, and the Ca^2+^ amplitude from the maximum and minimum signal across the plateau event as follows: 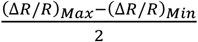. These datapoints were plotted in GraphPad Prism. To sort by derivative level (magnitudes), the ratiometric signal was sorted according to the derivative, then a cutoff was established where signals with a derivative lower than the median value were assigned to the intermediate derivative bin, while those with a derivative higher than the median were placed in the high derivatives bin. The data points were plotted in GraphPad Prism, where a Mann-Whitney test was used to determine no statistically significant differences between the 2 derivative levels.

#### Correlating the Lateral Head Bend and AFD Activity Oscillations

To isolate the high-frequency lateral undulations from the animal’s macroscopic forward trajectory, spatial coordinates during identified plateau events were mathematically rotated. First, linear regression was applied to the normalized spatial coordinates (*Y* vs. *X*_*norm*_) to determine the mean heading angle (*θ*). A rotation matrix was then constructed to rotate the coordinate system by *γ* (where *γ* = 90°−*θ*), effectively aligning the animal’s mean forward translation along the longitudinal axis. This orthogonal transformation isolated the spatial amplitude of the lateral head sweeps entirely onto a single transverse axis (*X*′_*vert*_). Prior to feature extraction, both the rotated positional coordinates and the empirical AFD calcium traces 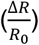 were optionally smoothed using a 1D Gaussian filter to minimize high-frequency tracking noise. To quantify and compare the rhythmic properties of the animal’s physical undulations and the corresponding AFD calcium dynamics, both the rotated spatial trajectory (*X*′_*vert*_) and the smoothed calcium trace were modeled as continuous sinusoidal functions over relative time (*t*): *f*(*t*) = *y*_0_ +*Asin* sin(*ωt* +*φ*), where *y*_0_ represents the baseline offset, *A* represents the oscillation amplitude, *ω* represents the angular frequency (in radians per second), and *φ* represents the phase shift.

Optimal model parameters were estimated via non-linear least squares optimization using the curve_fit function from the “scipy.optimize”Python library (See the Data and Code Availability section for the repository link). To ensure biological and mathematical plausibility, the initial estimates for angular frequency (*ω*_0_) were seeded based on an assumption of approximately 7 to 8 full periods per plateau event, and the optimization was executed utilizing the Trust Region Reflective (trf) algorithm. Parameter boundaries were strictly defined to restrict amplitudes to non-negative values (*A* ≥0) and to constrain angular frequency and phase within ±2*π*. The resulting angular frequencies were converted to standard frequencies 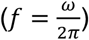 in Hertz (Hz) to allow direct phase and frequency comparisons between the mechanical motor output and the AFD calcium response. See the Data and Code Availability section for the repository link.

#### Quantification of Immobilized Animal Calcium Dynamics

Either a custom interpolator, or the TrackMate plugin in FIJI (Ershov et al., 2022) with manual identification and semiautomatic tracking were used to ensure the ROI covered the AFD soma at all time frames of the experiment’s time stack and returned the average intensities. This quantification was run separately for GCaMP6s and RFP. A custom Python script (See the Data and Code Availability section for the repository link) compiled fluorescence intensity in both channels across animals and experiments, aligned this information to temperature via the time stamps and calculated the ratio of the green GCaMP signal and RFP, and normalized with respect to the global minimum ratio (ΔR/R = (R_t_ - R_t-1_)/R_t-1_). To avoid overrepresenting each animal, regardless of successful dorsoventral positioning, only 1 AFD neuron was analyzed per worm.

### Fitting Responses Across Varying Thermal Ramps

To evaluate how the AFD neuron encodes temperature changes, the peak calcium response (maximum Δ*R*/*R*) was plotted against the temperature derivative across three paradigms: varying Δ*T*, varying Δ*t*, and thermal steps. The overall relationship across the full stimulus range was assessed using the Pearson correlation coefficient (*r*). The data was modeled using two regression strategies over defined stimulus windows. First, a linear fit was done. Peak responses within 0.1 to 0.6 °C/min were fit using ordinary least squares regression *y* = *αx* +*β*, where *y* is the peak Δ*R*/*R*, and *x* is the temperature derivative. All peak responses were then fit to a logistic sigmoid function. Empirical responses were first normalized to their maximum value and logit-transformed to allow for linear estimation of the growth rate (*α*) and horizontal offset (*β*). The model was then back transformed and scaled to the maximum empirical response to yield the final continuous curve 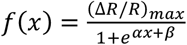. See the Data and Code Availability section for the repository link harboring the code used.

#### Deriving and Fitting the Response Function

First, the instantaneous temperature derivative was calculated for each experiment. Raw temperature and time arrays were downsampled by a factor of 10 (retaining every 10th data point) to dampen high frequency noise. The first derivative of these downsampled arrays was taken to calculate the change per second, and the resulting derivatives were subsequently linearly interpolated back to match the original array length and sampling frequency of the (Δ*R*/*R*).

The response function was then derived using a GLM solved via ordinary least squares regression. The stimulus matrix was constructed utilizing a rolling history window consisting of the prior 1,250 data points of the temperature derivative preceding each Ca^2+^ response data point, sampled at interval steps of 10. To isolate the neuron’s specific response to heating and remove negligible thermal noise, the stimulus array was rectified. Positive temperature derivatives strictly greater than 10^−7^ were retained, while negative derivatives (cooling) and near-zero fluctuations were set to exactly 0.0. The resulting least-squares solution vector yielded the temporal response function.

To establish a standardized (normalized) temporal filter across multiple experimental replicates, individual temporal response functions were linearly interpolated onto a common temporal grid with a fixed 0.3-second resolution. These interpolated functions were then averaged to compute a mean temporal response function. Finally, the decay kinetics of this mean response function were quantified by fitting a single exponential decay model defined by the equation: *f*(*t*) = *Ae*^−*Bt*^, where *A* represents the initial amplitude and *B* represents the decay rate constant. The parameters were estimated using non-linear least squares optimization. To ensure biological plausibility, the optimization was strictly bound to non-negative values for both amplitude and decay rate (*A* ≥0, *B* ≥0). See the Data and Code Availability section for the repository link harboring the code used.

#### AFD Calcium Dynamics Model

AFD calcium imaging datasets were analyzed across multiple thermal stimulus paradigms including a) freely moving worm navigating shallow or steep temperature gradients (Fig. 1), b) realistic temperature stimuli (Fig. 5), and c) sinusoidal up ramp protocols (Fig. 3). Raw temperature measurements were smoothed to reduce measurement noise while preserving stimulus dynamics using a Savitzky-Golay filter with polynomial order set to 2 and smoothing window of 3 seconds (Savitzky and Golay, 1964). The smoothed temperature trace was used for all downstream modeling. GCaMP6s can resolve fast calcium transient with an in vivo rise time of ~150 ms and half-decay time ~600 ms (Chen et al., 2013), more than an order of magnitude faster than the integration timescales captured by our model. GCaMP kinetics were therefore treated as negligible and not incorporated into the model.

Guided by our experimental findings, we modeled AFD calcium activity as arising from leaky integration of temperature derivatives. Specifically, we compute the instantaneous rate of temperature change, 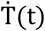, which is passed through a first-order leaky integrator to generate a latent dynamic state, D(t). The filtered derivative, D(t), was multiplicatively gated by an adaptive gain gamma window W(T(t)). By combining derivative detection, leaky integration, and temperature-dependent gain control, our mathematical model captures the core temperature encoding principles of AFD. In this formulation, the gain window is static within a given condition, with its parameters inferred from calcium data rather than dynamically adapted during an experiment. To determine the contribution of various components of the model, we developed and evaluated two nested conceptual models:

Model A: Temperature derivative followed by leaky integration

Model B: Output of Model A gated by the adaptive gain window

AFD calcium responses were modeled as a function of temperature dynamics through a sequence of transformations. The experienced temperature T(t) was first converted into its temporal derivative, 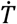. This signal was passed through a first-order leaky integrator to generate a latent drive variable D(t): 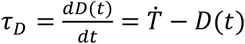, where *τ*_*D*_ is the time constant for leaky integration. In the baseline model (Model A), the predicted calcium response was obtained via a linear transformation: *F*_*pred*_(*t*) = *α* ·*D*(*t*) +*b*, where a and b are linear scaling parameter fits per animal. In the extended model (Model B), the drive signal was multiplicatively modulated by a temperature-dependent gain function *W*[*T*(*t*)]:

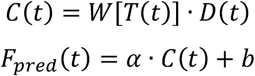

The gain function *W*[*T*(*t*)] was implemented as a normalized gamma probability distribution (PDF) evaluated at (*T* −*T*_*on*_), with values below *T*_*on*_ set to 0:

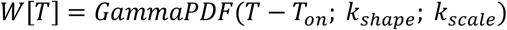

where *T*_*on*_, *k*_*shape*_ and *k*_*scale*_ determine the onset, shape, and width of the gain window, respectively. Model parameters were estimated independently for each animal. The integration timescale *τ*_*D*_ and gain parameters were inferred from experimental data, while a and b were obtained via least-squares regression.

To account for worm-to-worm variability in baseline and amplitude, for each model the output was mapped to calcium using a per-worm affine transformation. The coefficients of transformation were estimated by least-squares regression. Model parameters were optimized independently for each worm replicate using Optuna with a Tree-structured Parzen Estimator (TPE) sampler in Python (Akiba et al., 2019). Each model was fit using 300 trials per replicate. See the Data and Code Availability section for the repository link containing the code used.

#### Quantification of AFD sensitivity and Oscillation Prominence

Fluorescence data were analyzed ratiometrically to correct expression levels and movement artifacts. The fluorescence ratio *R* was calculated at each time point as 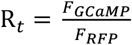. Baseline activity *R*_0_ was defined as the mean ratio averaged over 30s immediately preceding the stimulus at the initial temperature hold. Peak response *R*_*max*_ was determined as the maximum ratio observed during the stimulus period (temperature ramp). Neuronal sensitivity was quantified as the ratio fold change in fluorescence ratio relative to the baseline 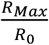. Plotted data are presented as individual values with overlaid mean ±standard deviation (SD). A Kruskal-Wallis with post-hoc Dunn’s test was conducted. Analysis was performed via a custom Python script (see Data and Code availability section for the repository link containing the code used).

Calcium oscillations were identified and quantified using the “scipy.signal”library in Python. Ratiometric traces *R* were first baseline-corrected 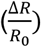 by subtracting the minimum value of the trace 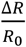 peaks were then identified as local maxima separated by a minimum inter-event interval of 15s across the entire experimental timeframe. Oscillation magnitude was defined by peak prominence, calculated as the vertical distance between the peak and the lowest contour line within a sliding window of 601 frames. Only peaks exceeding a minimum prominence threshold of 0.4 were included in the final analysis. Local sensory gain was estimated for each detected event by dividing the peak prominence by the mean positive temperature derivative calculated over the 10 frames (approximately 1 second) immediately preceding the peak. Analysis was performed via a custom Python script (see Data and Code availability section for the repository link containing the code used).

#### Analysis of Standard Thermotaxis Assays

From the acquired individual images, animals were identified and their coordinates over time obtained by using an adapted version of the MagatAnalyzer package previously described and custom MATLAB scripts also previously presented (Gershow et al., 2012; Hawk et al., 2018; Luo et al., 2014). The temperature coordinates derived from pixel position were then used to generate realistic temperature profiles that were delivered unto immobilized animals for calcium imaging.

To generate raster-style plots, x-coordinate data from each biological replicate were concatenated into a single .mat file. The combined dataset was then processed using rastergram_simpleinput.m, which converts positions into temperature values and bins the data according to the number of bins specified in the figure legend. For plotting population-level temperature experience over time, a time series for each biological replicate was computed by averaging temperature (converted from x-coordinates) within each time bin. The mean temperature and the associated standard error of the mean (SEM) were then calculated across replicates. The resulting data were plotted and formatted in Origin. See the Data and Code Availability section for the repository link harboring the codes used.

#### Quantification of run trajectories and run heading polar plots

Individual worm tracks were smoothed using a Savitzky-Golay filter (polynomial order 3, window width 7 frames) to reduce tracking noise while preserving curvature features. Within each segment, reorientation events were detected by computing the heading direction at each frame and identifying directional changes exceeding 120°within a sliding window. Run segments were defined as the intervals between two consecutive reorientation events, retaining only runs with a minimum duration of 5 seconds. For the run-trajectory plots, the top 40 longest runs per temperature region were selected, origin-shifted so each run began at the absolute cartesian origin coordinate (0,0). Two temperature bins were analyzed: Region 1 (20-22°C) and Region 2 (22-24°C), defined by the worm’s x-coordinate position mapped to temperature via the known linear temperature gradient. For the run-heading polar plots, run headings and total Euclidean run distances were computed for each run within each temperature region. Mean run distance was calculated per angular bin (bin width: 10°) and displayed as a polar plot with a fixed radial axis (0-6 mm). All analyses were implemented in MATLAB using custom. See the Data and Code Availability section for the repository link harboring the code used.

#### Simulation of thermotactic navigation driven by AFD encoding

To test whether the experimentally derived AFD encoding principles are sufficient to reproduce thermotactic behavior, we implemented a minimal simulation framework linking sensory input to locomotory decisions. Each simulated worm navigated a linear thermal gradient for up to 40 minutes (timestep = 1 s). The simulation is based on the AFD temperature encoding model derived in this paper (Model B).. At each timestep, the instantaneous temperature derivative was determined. It was then passed through a first-order leaky integrator to yield a latent-drive variable. This latent-drive variable was multiplied by a temperature-dependent gain, implemented as a gamma-shaped function to produce an effective behavioral drive. The parameters for the simulation such as time constant, gamma distribution constants were obtained from the AFD calcium model fits, while the locomotion parameters such as run durations, speed, etc., were obtained from the behavioral assays. At each run initiation, the probability of entering isothermal tracking mode was computed as a logistic function of the behavioral drive. Higher values of behavioral drive biased movement toward isothermal movement, while lower values promoted movement parallel to the thermal gradient. Simulations were run under the steep gradient assay configuration (15-25°C, 10 ×10 cm arena) with simulated worms starting temperatures of 20°C. The simulation was implemented in Python. See the Data and Code Availability section for the repository link harboring the code used.

#### Statistical Analysis

All statistical analysis was done via Python or GraphPad Prism, with Kruskal Wallis and Dunn’s post hoc test for analyzing multiple comparisons or Two-tailed Mann-Whitney for comparing 2 groups, unless otherwise stated.

#### Declaration of generative AI

During the preparation of this work, the authors used: 1) ChatGPT to assist with textual edits for clarity, and to assist in code modification; and 2) ResearchRabbit to help scan the existing literature, and to identify and assess relevant literature to the study. After using this tools/services, the authors reviewed and edited the content and take full responsibility for the content of the published article.

**Table 1.** Key reagents and strains.

| REAGENT or RESOURCE | SOURCE | IDENTIFIER |
| --- | --- | --- |
| Chemicals |  |  |
| Levamisole Hydrochloride | Sigma-Aldrich | SKU#PHR1798; |
|  |  | CAS: 16595-80-5 |
| **Experimental Models: Organisms/Strains** |  |  |
| N2 (wild type) | Caenorhabditis Genetics Center | N2 |
| *C. elegans: Wyls629 (pgcy-8::GCaMP6s)* | (Hawk et al., 2018) | DCR Lab Strain ID: DCR3055 |
| *C. elegans: Co-labeled Freely Moving Calcium Imaging: N2; olaEx5141[ (pgcy8::tagRFP::unc54)(30ng/μl) + (pgcy8::GCaMP6s::unc54)(30ng/μl)+ (pelt7::GFP::unc54)(30ng/μl) ]* | This paper | DCR Lab Strain ID: DCR8464 |
| *C. elegans: Co-labeled Freely Moving Calcium Imaging: N2; olaEx5146[ (pgcy8::tagRFP::unc54)(30ng/μl) + (pgcy8::GCaMP6s::unc54)(30ng/μl)+ (pelt7::GFP)(30ng/μl) ]* | This paper | DCR Lab Strain ID: DCR8483 |
| **Recombinant DNA** |  |  |
| Plasmid: pgcy8::tagRFP::unc54 | This paper | DCR Lab Strain ID: DACR1422 |
| Plasmid: pgcy8::GCaMP6s::unc54 | This paper | DCR Lab Strain ID: DACR943 |
| Plasmid: pelt7::GFP::unc54 | This paper | DCR Lab Strain ID: DACR2353 |

| Software and Algorithms |  |  |
| --- | --- | --- |
| MicroManager (Fiji) V.<br>1.4/2.0 | Virtual | https://download.micro-manager.org/release/1.4/Windows/MMSetup_32bit_1.4.22.exe |
| Fiji | (Schindelin et al., 2012) | https://imagej.net/Fiji/Downloads |
| LabView 2011 | National Instruments | www.ni.com/en.html?cid=PSEA-7013q000001fLKAAA2-CONS-Bing_1239149829979437&utm_keyword=national%20instruments&msclkid=ab7b918cdac01b423c37f716bfc82087 |
| TrackMate | (Ershov et al., 2022) | https://imagej.net/plugins/trackmate/ |
| BioRender | BioRender.com | https://app.biorender.com/biorender-templates |
| MATLAB | MathWorks | https://www.mathworks.com/products/matlab.html |
| Python 3.8.0/3.12.1 | Python Software Foundation | https://www.python.org/ |
| GraphPad Prism | Graphpad Software Inc | https://www.graphpad.com/ |
| ApE | M. Wayne Davis | https://jorgensen.biology.utah.edu/wayned/ape/ |

**Table 2.** Freely Moving Animals Imaging Deep Learning Processing Codes and GitHub Links.

| Code Name | Purpose | GitHub Link |
| --- | --- | --- |
| FIJI Merge Channels Script GCAMP6+RFP | FIJI macro code used to overlay dual channel recordings and | Here |
|  | prepare for behavior analysis. |  |
| DLC Config + Associated Files | DLC config code used to set the parameters of the convoluted neural network (CNN) for training. | Here |
| Freely Moving Analysis Example Code:<br>Full_Cal_Model-Copy1(3_4_20_17to24_1_2_).ipynb<br>DLC_Behavior_Analysis .ipynb | Code used to take DLC output, allowing for the construction of figures. | Main folder<br>Subfolder 1<br>Subfolder 2 |
| Neuron_Tracker_Codes | Code used to reliably record neuronal calcium activity as worms freely navigate the thermal gradient. | Here |
| FIJI_Split_Channels_GCaMP/RFP | ImageJ code used to split dual channels, in order to individually detect RFP and GCaMP6 fluorescent neurons. | Here |
| CytoSHOW > Ratio Chaser (megaTiff<br>MicroManager)<br>(ImageJ=>BETA) | CytoShow used to record and analyze calcium imaging data in freely moving worms performing thermotaxis. | Here |
| AFD_Model_DeepLearning | Files used to predict a binary AFD skeleton mask for every individual green and red fluorescent channel image, which was then used to precisely quantify the location and pixel intensity of the GCaMP and RFP in all parts of the neuron. | Here |

## Figure Legends

**Figure S1.**
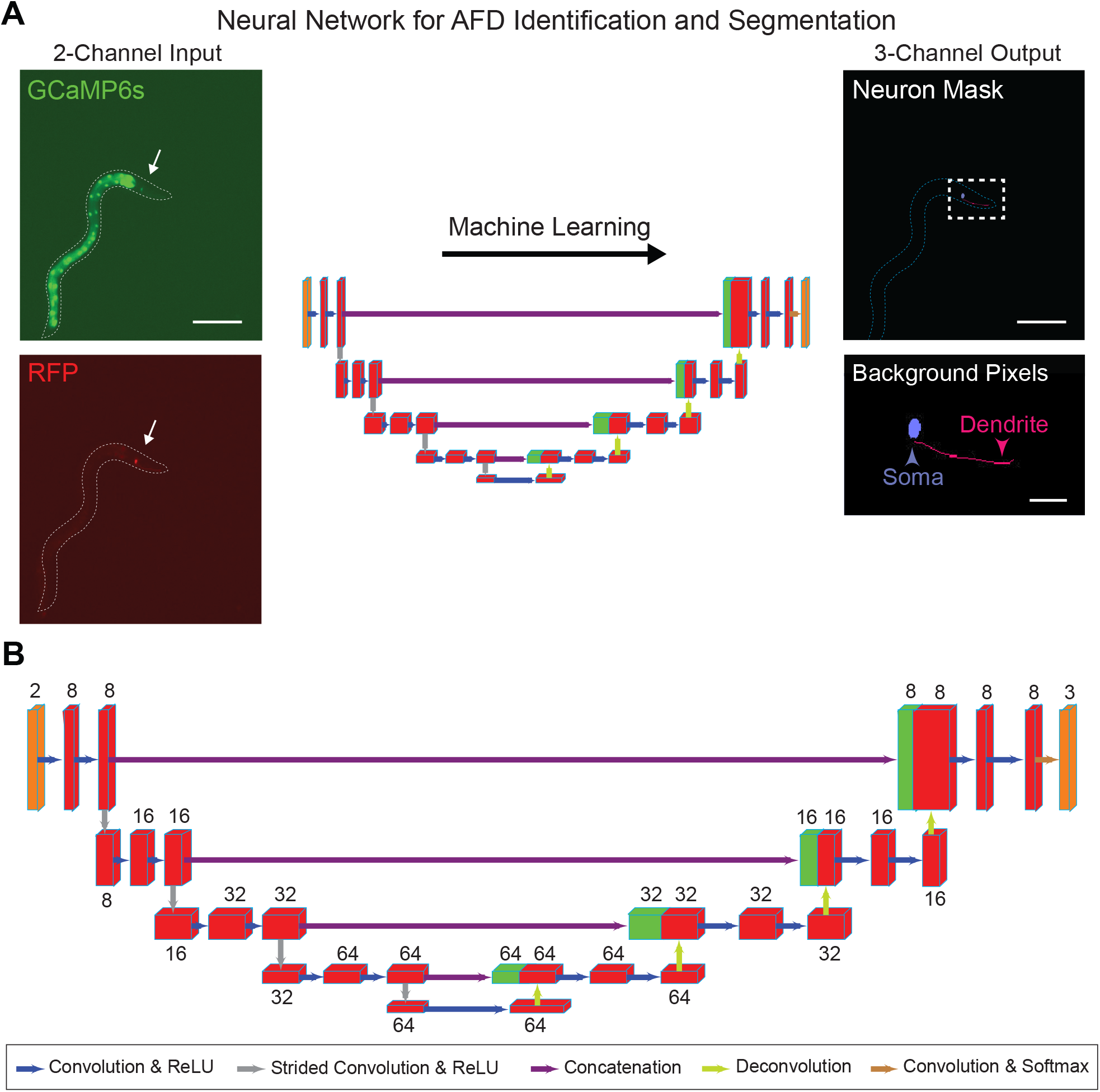
related to Figure 1. Analysis pipeline for AFD neuron detection and quantification using a 3D U-Net. **(A)** Example input and output images for the 3D U-Net, a type of convolutional neural network (CNN). Each input image is two-channel (red and green) with identical dimensions. The arrow points to the AFD neuron. A sequence of such images is provided to the CNN for processing. A post-CNN processing step creates a pixel skeleton of AFD that was used to extract the corresponding pixel coordinates for both the soma and the dendrite (right side). These XY coordinates of the neuron skeleton mask were used to extract the corresponding fluorescence intensity pixels in the input image. See the Methods section Deep Learning Image Processing of Freely Moving Animal Thermotaxis Data for more information. The neuron coordinates and fluorescent activity were then used for further analysis. (Left scale bar, 250 μm; top right scale bar, 250 μm; bottom right scale bar, 50 μm.) **(B)** Schematic of the 3D U-Net architecture comprising layers which processed the input images sequentially, going from left to right. Each layer transforms the preceding block, starting with a two-channel input image and finally outputting a three-channel probability map. The CNN was trained to distinguish between the soma and dendrite of the AFD neuron and background within each image. Each pixel in the output map reflects the likelihood of the pixel being part of the soma, the dendrite, or background. Numbers above each block denote the number of channels in the operation; a higher channel count requires more computation but is better suited to classify pixels accordingly. Red blocks represent outputs of convolution (blue arrows) and deconvolution (lime green arrows), whereas green blocks are concatenated to other blocks (purple arrows). Input and output blocks are depicted in orange.

**Figure S2.**
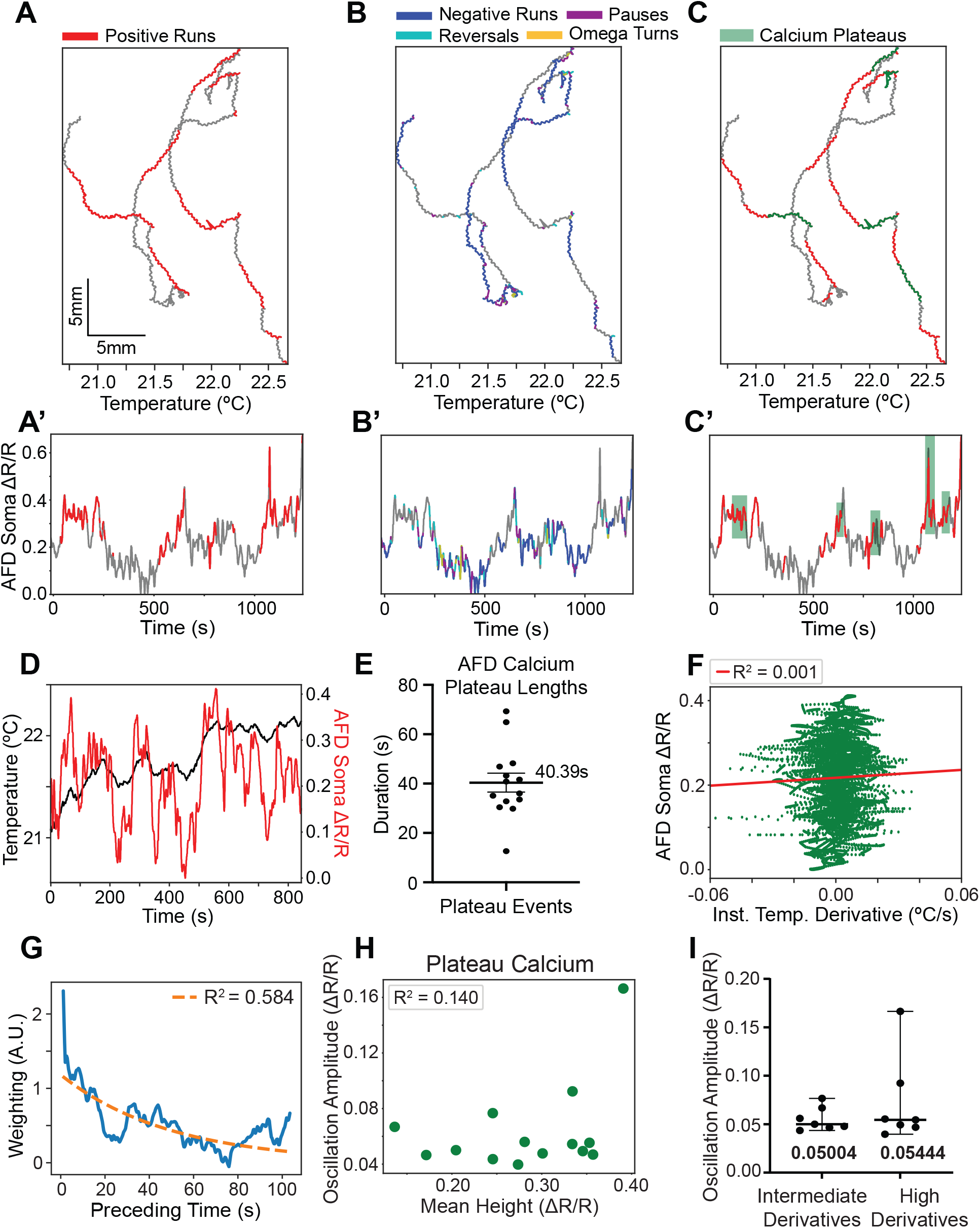
related to Figure 1. Additional analyses of AFD calcium dynamics and behavior in freely moving thermotaxis. **(A)** Additional representative thermotaxis trajectories of a 25°C-trained animal navigating a gradient. Positive runs colored red. **(B)** Same track shown in **(A)**, with colored annotations for other behavioral strategies (negative runs in blue, reversals in cyan, omega turns in yellow and pauses in purple). **(C)** Plateau events colored green on the same track as in **(A-B). (A’-C’)** Corresponding AFD soma Ca^2+^ (ΔR/R) traces for the tracks shown in the top panels. **(D)** AFD soma Ca^2+^ (ΔR/R) (red) overlaid on the temperature experienced by the animal (black line) over the course of the experiment shown in Fig. 1B-F. **(E)** Mean duration of labeled AFD Ca^2+^ plateaus. Each point (n=14 from 3 independent experiments) denotes the length of 1 plateau and error bars denote SEM. **(F)** Instantaneous temperature derivative and corresponding AFD soma Ca^2+^ across all time points for the same experiment shown in Fig. 1. Red line shows linear relationship (R^2^=0.001). **(G)** Impulse-response (weighting) function derived from freely moving data (n=3 experiments), illustrating that recent temperature derivatives contribute most strongly to the present Ca^2+^ level, with generally smaller weights assigned to more distant time points, consistent with (Tsukada et al., 2016). The time constant of the best fit one phase decay curve is τ= 49.9s. See the Methods section Deriving and Fitting the Response Function. **(H)** Relationship between amplitude of oscillations in AFD Ca^2+^ plateaus and the mean height of the plateau calcium level (ΔR/R) in freely moving animals. Each point (n=14) represents 1 plateau event from 3 independent experiments. **(I)** Median amplitude of oscillations in AFD Ca^2+^ plateaus and the corresponding temperature derivative experienced by animals during each plateau, binned by intermediate and high derivative levels. Bins were established by sorting derivatives by magnitude and using the median as cutoff. Each point (n=7 per bin) denotes a plateau across 3 independent experiments and error bars denote 95%confidence interval. Two-tailed Mann-Whitney test showed no statistically significant differences.

**Figure S3.**
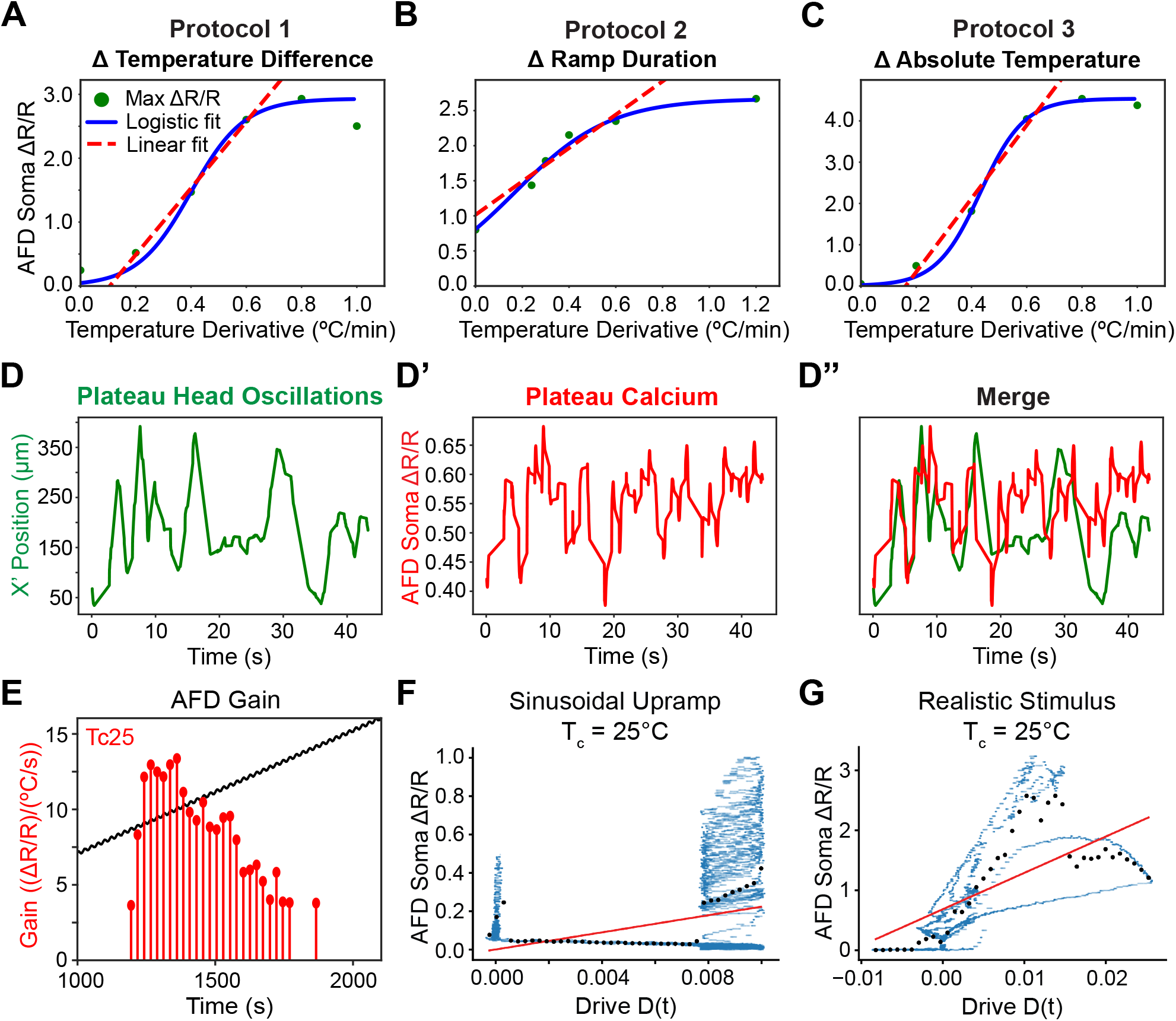
related to Figures 2, 3, and 5. Relationship of Ca^2+^ levels to experienced temperature derivatives. **(A)** AFD response versus temperature derivative for the protocol of Fig. 2A. Each point corresponds to the peak of the mean response obtained for a single ramp experiment, and both linear and logistic fits were applied. **(B)** As in **(A)**, but for Fig. 2B. **(C)** As in **(A-B)**, but for Fig. 2C. **(D-D″)** Representative segment from a Ca^2+^ plateau event demonstrating the relationship between head position on a thermal gradient and AFD activity. **(D)** Adjusted AFD soma x-position over time, with oscillations isolated from forward drift movement by projecting coordinates over X’axis. **(D′)** Simultaneously recorded AFD soma Ca^2+^ (ΔR/R) from the same time window. **(D″)** Overlay of adjusted x-position and AFD Ca^2+^, illustrating their correspondence. **E**. Local sensory gain of AFD oscillations from the Tc25 animal response highlighted in Fig. 3B. Individual markers and stems represent the estimated gain for each detected calcium event. Gain is calculated as the ratio of peak prominence to the mean positive temperature derivative 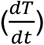 **(F-G)** Filtered derivative drive-calcium relationship does not follow a linear relationship. Binned average measured Ca^2+^ (ΔR/R) against model derived D(t). Black dots denote binned means of the time points in blue, and the red line a linear fit. **(F)** For sinusoidal upramp, as in Fig. 5F. **(G)** For realistic stimulus, as in Fig. 5G.

